# Bacteria contribute to plant secondary compound degradation in a generalist herbivore system

**DOI:** 10.1101/865212

**Authors:** Charlotte B. Francoeur, Lily Khadempour, Rolando D. Moreira-Soto, Kirsten Gotting, Adam J. Book, Adrián A. Pinto-Tomás, Ken Keefover-Ring, Cameron R. Currie

## Abstract

Herbivores must overcome a variety of plant defenses, including coping with plant secondary compounds (PSCs). To help detoxify these defensive chemicals, several insect herbivores are known to harbor gut microbiota with the metabolic capacity to degrade PSCs. Leaf-cutter ants are generalist herbivores, obtaining sustenance from specialized fungus gardens that act as external digestive systems, degrading the diverse collection of plants foraged by the ants. There is *in vitro* evidence that certain PSCs harm *Leucoagaricus gongylophorus*, the fungal cultivar of leaf-cutter ants, suggesting a role for the Proteobacteria-dominant bacterial community present within fungus gardens. Here, we investigate the ability of symbiotic bacteria present within fungus gardens of leaf-cutter ants to degrade PSCs. We cultured fungus garden bacteria, sequenced the genomes of 42 isolates, and identified genes involved in PSC degradation, including genes encoding cytochrome p450s and genes in geraniol, cumate, cinnamate, and α-pinene/limonene degradation pathways. Using metatranscriptomic analysis, we show that some of these degradation genes are expressed *in situ*. Most of the bacterial isolates grew unhindered in the presence of PSCs and, using GC-MS, we determined that isolates from the genera *Bacillus, Burkholderia, Enterobacter, Klebsiella,* and *Pseudomonas* degrade α-pinene, β-caryophyllene, or linalool. Using a headspace sampler, we show that sub-colonies of fungus gardens reduced α-pinene and linalool over a 36-hour period, while *L. gongylophorus* strains alone only reduced linalool. Overall, our results reveal that the bacterial community in fungus gardens play a pivotal role in alleviating the effect of PSCs on the leaf-cutter ant system.

**Importance:** Leaf-cutter ants are dominant neotropical herbivores capable of deriving energy from a wide range of plant substrates. The success of leaf-cutter ants is largely due to their external gut composed of key microbial symbionts, specifically, the fungal mutualist *L. gongylophorus* and a consistent bacterial community. Both symbionts are known to have critical roles in extracting energy from plant material, yet comparatively little is known about their role in the detoxification of plant secondary compounds. Here, we assess if the bacterial community associated with leaf-cutter ant fungus gardens can degrade harmful plant chemicals. We identify plant secondary compound detoxification in leaf-cutter ant gardens as a process that depends on the degradative potential of both the bacterial community and *L. gongylophorus*. Our findings suggest the fungus garden and its associated microbial community influences the generalist foraging abilities of the ants, underscoring the importance of microbial symbionts in plant substrate suitability for herbivores.

## Introduction

Plants defend themselves from herbivores through the production of an extraordinarily diverse set of plant secondary compounds (PSCs) (1–4). Not only are these chemicals diverse in structure and toxicity, the mechanisms behind their synthesis, storage, and release are complex, depending on the type of damage, the taxon inflicting the damage, the species of plant, environmental conditions (e.g., hours of sunlight, temperature, moisture), and other various factors (3, 5, 6). The range of plant chemical defenses require herbivorous insects to engage in multiple strategies to counter toxic effects of PSCs, including harboring microbial symbionts that aid in detoxification. Microbes associated with mountain pine beetles (7–9), red turpentine beetles (8), pine weevils (10), gypsy moths (11), apple maggot flies (12), termites (13), and coffee berry borers (14) have been found to play an important role in PSC degradation. Understanding the role of microbial symbionts in PSC detoxification is critical, since the capacity of insects to mitigate PSC toxicity is an important factor in determining host plant range (15).

Leaf-cutter ants, two genera within the monophyletic fungus-growing ant subtribe, are dominant herbivores in most Neotropical ecosystems and are able to forage from a diverse array of plants. In a long-term study, active colonies of two species of leaf-cutter ants, *Atta colombica* and *Atta cephalotes*, were observed cutting leaves from 67–77% of all plant species (86 and 59 plant species, respectively) recorded in their respective foraging areas (16). In a year-long study by Wirth and colleagues (17), one colony of *A. colombica* foraged from the leaves or flowers of 126 species, representing 91 genera and 52 families. In mature colonies of *Atta* sp., the ants forage ravenously, forming massive foraging columns that create distinctive trails (Figure 1A). The leaf-cutter ants do not directly consume the plant substrate, but rather use it to feed *Leucoagaricus gongylophorus* (Agaricales: Agaricaceae), the obligate fungal mutualist. In return, *L. gongylophorus* degrades the leaf substrate and serves as food for the ants, providing energy and nutrients in the form of specialized hyphal swellings known as gongylidia (18, 19). This process occurs in structures known as fungus gardens (Figure 1B), which are maintained in underground chambers. Other lineages of fungus-farming ants bring non-leaf substrate to *Leucoagaricus* sp. and contain 1-20 fungus garden chambers with a total colony size of no more than a few thousand workers (20, 21). In contrast, mature colonies of the leaf-cutting ant genus *Atta* can be composed of hundreds of fungus garden chambers, which provide nutrition to millions of larvae, pupae, and emerged workers (19, 22).

**Figure 1.**
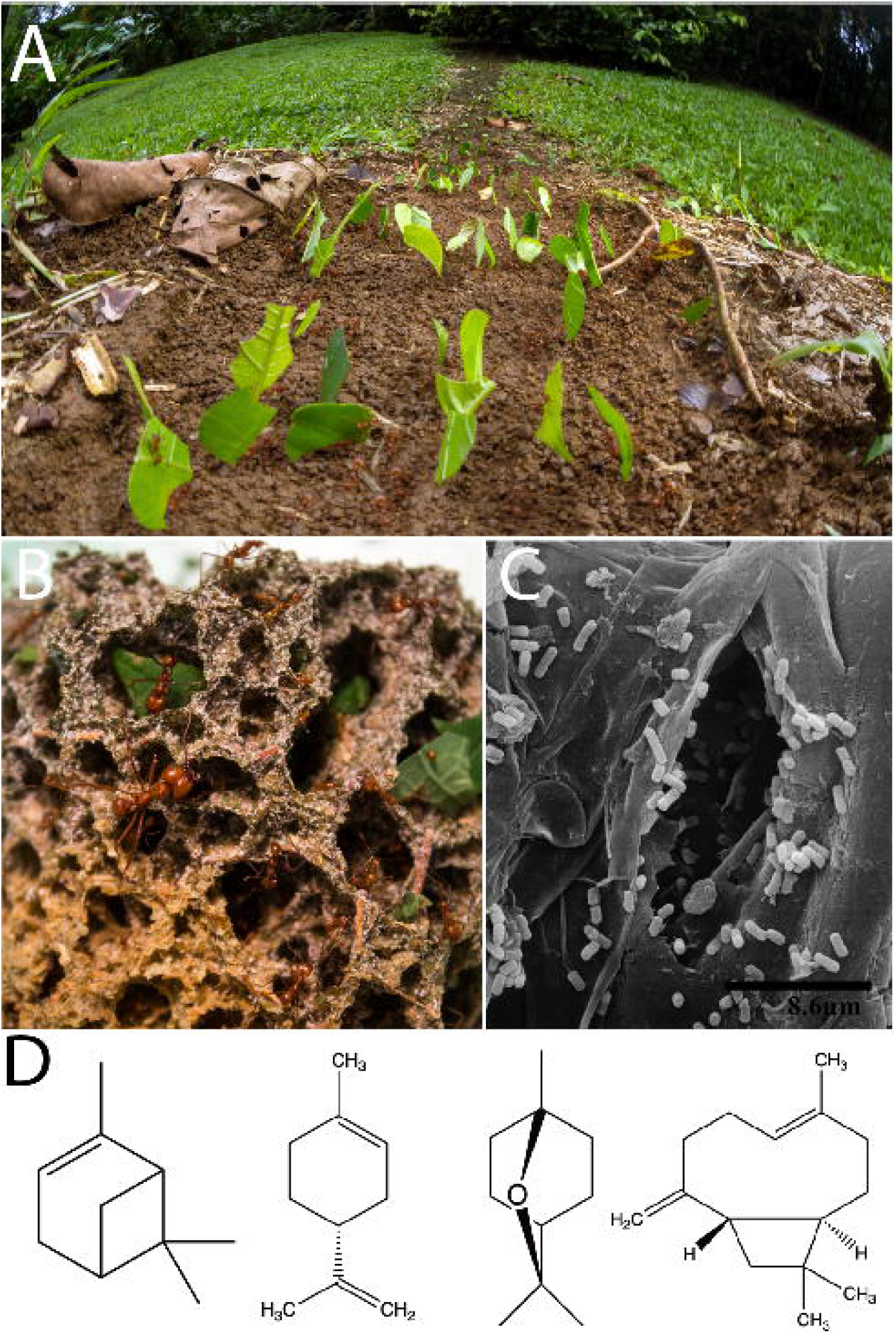
Leaf-cutter ants are dominant herbivores through the formation of multiple microbial symbioses. Leaf-cutter ants (*Atta cephalotes*) cut fresh leaf material at large scales, forming visible foraging trails (A) to bring the leaf material as a food source to their specialized fungus gardens (B). Bacteria are known to co-exist within the fungus garden, as seen in this SEM image (C). We can detect PSC from the fresh leaf material in the fungus gardens using GC-MS. One *Atta laevigata* sample collected in an area with eucalyptus had detectable amounts of PSC, such as α-pinene, *p*-cymene, eucalyptol, and caryophyllene oxide (31) (D). Photo credits: panel A Alexander Wild; panel B Lily Khadempour; panel C Rolando Moreira-Soto, reprinted from reference (25).

The fungus garden, which functionally serves as the ants’ external gut, includes the main fungal symbiont *L. gongylophorus* and a diverse and abundant community of bacteria. In contrast, the internal gut of leaf-cutter ants has a reduced bacterial community, with adult worker guts associated with *Wolbachia* or Mollicutes (23, 24). Culturing, scanning electron microscopy (25) (Figure 1C), and metagenomics of fungus gardens demonstrate a consistent presence of garden bacteria and have established a core bacterial community that consists mostly of Proteobacteria, the majority being in the class Gamma-Proteobacteria (26–31). Metagenomic studies show evidence for potential roles of garden bacteria in amino acid, vitamin, iron, and terpenoid nutrient supplementation, among other processes (26, 31). However, metagenomic predictions cannot establish these roles definitively, as they only demonstrate that garden bacteria have the genetic potential to carry out these functions. Attempts have been made to use proteomics (32) and transcriptomics (33) to examine the bacterial community *in situ*, however, this has not proven fruitful due to the low ratio of bacterial to fungal biomass in fungus gardens. Fortunately, easily culturable bacterial genera, such as *Burkholderia, Enterobacter, Klebsiella, Pantoea,* and *Pseudomonas* are known to be consistently present in fungus gardens. The advantage of having cultured isolates was leveraged in a previous study with *Klebsiella* and *Pantoea,* demonstrating the critical role of these two genera in fixing nitrogen that assimilates in the bodies of the leaf-cutter ants (34).

In this study, we examined the ability of garden bacteria from leaf-cutter ants to metabolize PSCs. We focus on bacteria isolated from fungus gardens of fungus-growing ants to investigate the potential of garden bacteria to tolerate and degrade PSC. First, we used previously-isolated strains of *L. gongylophorus* and *Leucoagaricus* sp. and determined their susceptibility to eight PSCs. Next, we sequenced the genomes of 42 isolates of garden bacteria and predicted the presence of genes involved in PSC degradation. We exposed garden bacteria isolates to eight PSCs and determined their susceptibility. Then, using gas chromatography-mass spectrometry (GC-MS), we quantified the *in vitro* ability of 15 isolates of garden bacteria to degrade four PSCs. Finally, we measured reduction of two PSCs by fungus gardens from our laboratory colonies of *Atta cephalotes* using a headspace sampler coupled to a gas chromatograph. For additional evidence, we analyzed previously generated garden bacteria metagenomes, *Leucoagaricus* sp. genomes, and metatranscriptomes to investigate the presence and expression of genes involved in PSC degradation.

## Results

### *L. gongylophorus* and *Leucoagaricus* sp. tolerance of PSC

Four different strains of *L. gongylophorus* and one strain of *Leucoagaricus* sp. were tested for their ability to grow in the presence of eight different PSCs (Figure 2A-B). The chosen PSCs are likely to be encountered by leaf-cutter ants based on foraging studies. The *Leucoagaricus* sp. from *Paratrachymyrmex diversus* was the most generally inhibited, with complete growth inhibition from terpinolene, eucalyptol, linalool, and *p*-cymene, and high inhibition from α-pinene and limonene. The *Leucoagaricus* sp. from a *P. diversus* colony and *L. gongylophorus* from an *Atta laevigata* colony were also the only fungal cultivars that were inhibited by the sesquiterpene β-caryophyllene. The *L. gongylophorus* from an *Atta sexdens* colony also exhibited high sensitivity to PSC, with complete inhibition occurring from limonene, terpinolene, eucalyptol, and linalool. The *L. gongylophorus* from an *Atta capiguara* colony was the most resistant to the PSC tested, only exhibiting high inhibition in the presence of linalool, while exhibiting low-to-no inhibition in the presence of the remaining seven compounds (Figure 2B).

**Figure 2.**
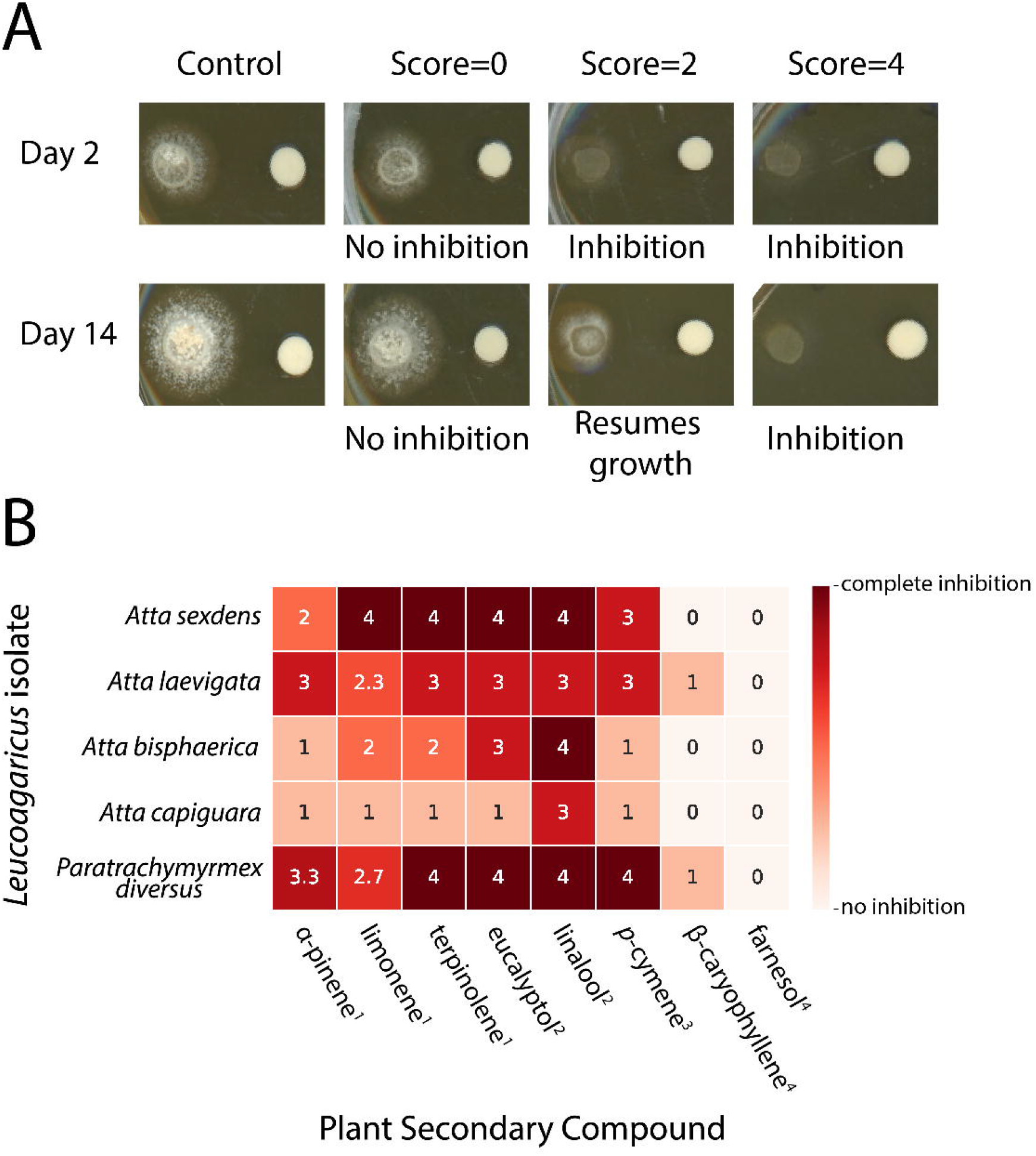
*L. gongylophorus* and *Leucoagaricus* sp. strains have variable tolerance of different PSC. The growth of *L. gongylophorus* and *Leucoagaricus* sp. was qualitatively scaled (A). We exposed each isolate to eight PSCs using a disc assay, done in triplicate. We then scored the growth for each isolate and averaged across the 3 technical replicates (B). The terpene class is indicated by the superscript number. 1= monoterpene, 2=terpenoid, 3= alkylbenzene related to monoterpene, 4=sesquiterpene.

#### Gene content of bacterial isolates, garden bacteria metagenomes, *L. gongylophorus*, and *Leucoagaricus* sp

We conducted BLAST-based and Kyoto Encyclopedia of Genes and Genomes (KEGG) annotations on whole genomes of 42 bacteria isolated from fungus gardens, garden bacteria metagenomes (31), one *L. gongylophorus* genome, and one *Leucoagaricus* sp. genome to assess the presence of genes involved in PSC degradation. No genes belonging to the diterpene degradation cluster, the cymene pathway, or *saxA* were found in the 42 bacterial isolates analyzed (Dataset S1, Dataset S2). However, the metagenome annotation did predict the presence of the cymene degradation pathway: one metagenome had the complete pathway, two metagenomes had ¾ of the pathway, five metagenomes had one gene from the pathway, and the remaining four had none of the genes (Dataset S2).

The α-pinene/limonene degradation pathway and geraniol pathway shared three genes between the two pathways: K01692 (enoyl-CoA hydratase), K01825 (fadB, 3-hydroxyacyl-CoA dehydrogenase), and K01782 (fadJ, 3-hydroxyacyl-CoA dehydrogenase). All isolates had at least one out of three shared genes, while *Acinetobacter, Enterobacter, Klebsiella,* and one *Pantoea* had all three. For the unique genes in the pathways (9 genes out of 12 for α-pinene/limonene and 12 genes out of 15 for geraniol), four of seven *Enterobacter* isolates had the highest proportion of unique α-pinene/limonene genes at 22%. All *Burkholderia* isolates, the *Acinetobacter* isolate, four of seven *Enterobacter* isolates, one of four *Klebsiella* isolates, and one of ten *Pantoea* isolates were predicted to contain the gene encoding monoterpene epsilon-lactone hydrolase (K14731), which is involved in monocyclic monoterpene degradation (35). Most isolates also contained a gene encoding aldehyde dehydrogenase (K00128) involved in the α-pinene/limonene degradation pathway (Figure 3A). In the 12 metagenomes, the proportion of genes present from the α-pinene/limonene pathway ranged from 22% to 56% (Dataset S2).

**Figure 3.**
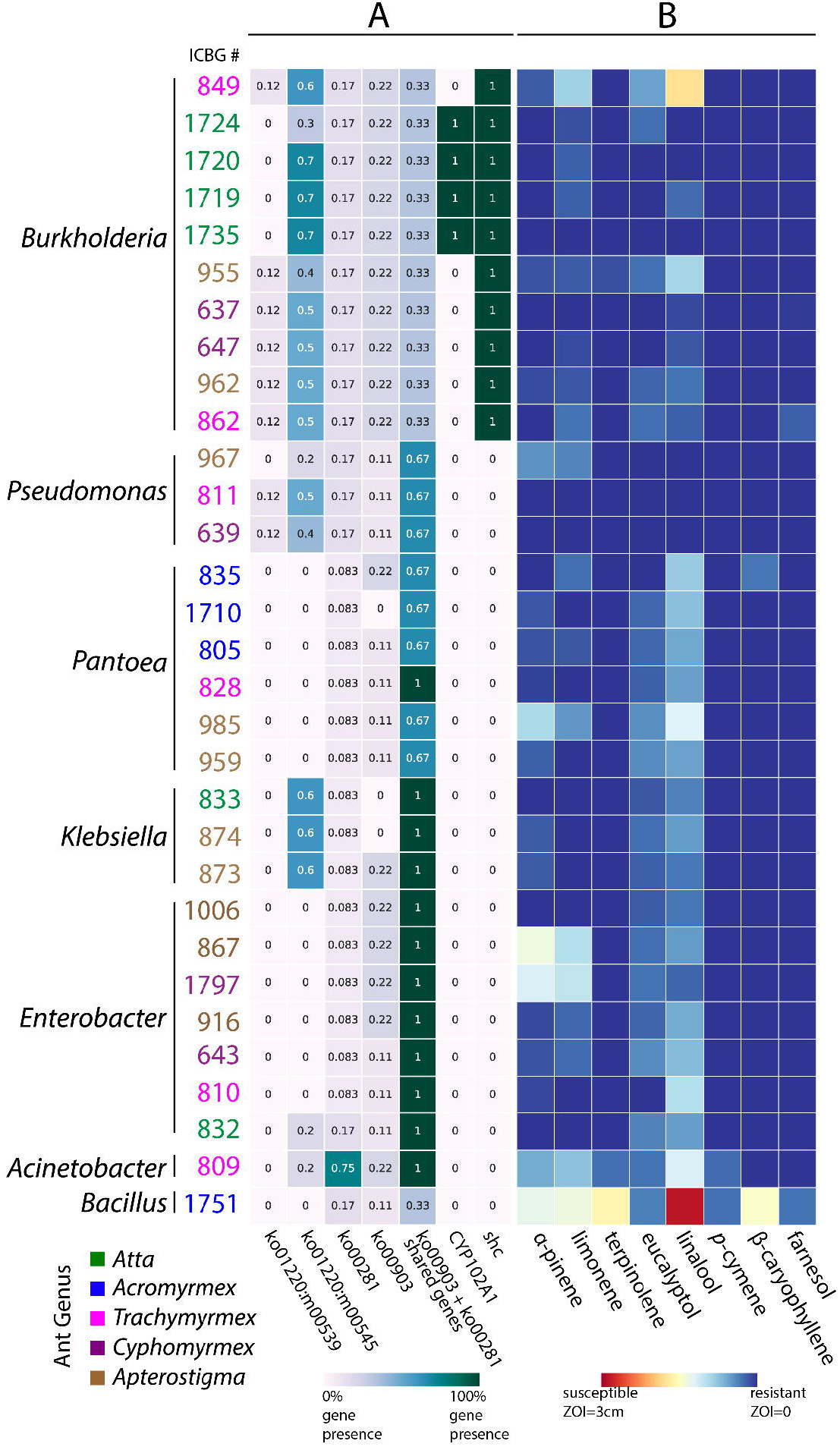
Gene content and PSC tolerance of fungus garden bacteria. Bacteria isolated from different genera of fungus-farming ants (isolate numbers color-coded by colony of origin) are predicted to have genes involved in PSC transformation (A) and are able to survive in the presence of eight PSC (B). (A) ko01220:m00539 corresponds to the cumate degradation module in the degradation of aromatic compounds pathway. ko01220:m00545 corresponds to the trans-cinnamate degradation module in the degradation of aromatic compounds pathway. ko00281 corresponds to the geraniol degradation pathway. ko00903 corresponds to the α-pinene/limonene degradation pathway. The shared genes between the geraniol pathway and the α-pinene/limonene pathway are K01692, K01825, and K01782. All of the pathways are presented as proportions with 1 being 100% of the genes predicted to be present and 0 being 0% of the genes predicted to be present. CYP102A1 and shc are single genes and so 0 or 1 indicates the absence or presence of that sequence, respectively. 31/42 annotations are displayed here, the annotations for the remaining 11 genomes are included in supplementary, in addition to the metagenome and metatranscriptome annotations (Tables S3, S4, S7) (B) We tested 46 isolates but 31 isolates are displayed as these are isolates that have genomes and were tested for tolerance. Data from all 46 isolates are provided in Figure S1.

For the unique genes in the geraniol pathway, the *Acinetobacter* isolate had the highest proportion of genes at 75%, while most other isolates were predicted to have between 8% and 17% of genes (Figure 3A). *Acinetobacter* contained all the genes except for the genes encoding geraniol dehydrogenase (K19653, K17832) and one of the 3-hydroxyacyl-CoA dehydrogenases (K00022). The other isolates were predicted to have genes encoding hydroxymethylglutaryl-CoA lyase and acetyl-CoA acyltransferase, in addition to the shared genes described above. In the metagenomic dataset, the proportion of completeness for the geraniol degradation pathway was between 17% and 75% (Dataset S2).

Ten of the isolates contained one gene from the cumate pathway, with seven *Burkholderia* and two *Pseudomonas* isolates predicted to contain genes encoding different components of the p-cumate 2,3-dioxygenase enzyme (K16303 or K16304) and one *Pantoea* isolate predicted to have a gene encoding 2,3-dihydroxy-p-cumate/2,3-dihydroxybenzoate 3,4-dioxygenase (K10621). On the other hand, more isolates had genes involved in the trans-cinnamate transformation pathway. Ten of eleven *Burkholderia*, all *Klebsiella*, and one of three *Pseudomonas* isolates had at least 50% of the genes necessary for cinnamate degradation (Figure 3A). The one *Paraburkholderia* isolate is predicted to contain 80% of the cinnamate pathway. Not all of the genes are necessary for a complete pathway, as there are two routes from trans-cinnamate to trans-2,3-dihydroxy-cinnamate (KEGG module M00545). In the metagenomes, cumate and trans-cinnamate pathways had a range of 0%-100% and 50%−80% completeness, respectively (Dataset S2).

Twenty cytochrome p450s known to be involved in isoprenoid transformation were analyzed (36). Most of the cytochrome p450s were not detected in the 42 bacterial isolates tested (Dataset S1). However, four cytochrome p450s encoding genes were predicted to be present in these genomes: CYP102A1, CYP106A2, CYP107H, and CYP108. CYP102A1 was detected in four out of 11 *Burkholderia* isolates, but none of the other genera (Figure 3A), CYP106A2 was detected in *Bacillus* (ICBG1751), and CYP107H and CYP108 were both detected in one *Pantoea* isolate (ICBG870). We saw an increase in the amount of genes encoding cytochrome p450s predicted to be present in the metagenomes, including CYP111, which catalyzes the 8-methyl hydroxylation of linalool (37). The other cytochrome p450 encoding genes detected only in the metagenomes were CYP105A3, CYP101, CYP105A1, as well as the cytochrome p450s described above in the individual bacterial genomes: CYP102A1, CYP107H, CYP106A2, and CYP108 (Dataset S1).

All *Burkholderia* isolates and the *Asaia* isolate genomes were predicted to encode a squalene-hopene cyclase (*shc*), while none of the other isolates were predicted to contain this gene (Figure 3A). The gene encoding *shc* was detected in 11/12 metagenomes (Dataset S1).

In addition to the garden bacteria genomes and metagenomes, we analyzed two existing fungal cultivar genomes for the same PSC degradation pathways. Overall, the two genomes, one from *A. cephalotes* (*L. gongylophorus* [Ac12]) and one from *Cyphomyrmex costatus* (*Leucoagaricus* sp. [SymC.cos]), lacked most of the gene sets analyzed (Dataset S1, Dataset S2).

##### Bacterial tolerance of PSC

Most bacterial isolates were able to grow uninhibited in the presence of the eight different compounds (Figure 3B). Linalool was the most inhibitory against the bacterial isolates, causing some degree of inhibition against all isolates, except *Pseudomonas* and three *Burkholderia* isolates. Farnesol, β-caryophyllene, and terpinolene did not inhibit the majority of bacterial isolates, causing small zones of inhibition in the *Bacillus* isolate as well as one to two other isolates in the genera *Burkholderia* and *Pantoea*. α-pinene and limonene caused slightly more inhibition, especially in *Bacillus* and two *Klebsiella* isolates, but most bacteria were resistant or only slightly susceptible to these two compounds. These trends were also seen in a larger set of samples (Figure S1).

##### GC-MS of bacterial isolates incubated with PSC

All *Enterobacter* isolates (ICBG810, *P*=0.0004; ICBG643, *P*=0.0018; ICBG832, *P*=0.0062) significantly reduced α-pinene (*t*-test, Bonferroni correction: α=0.0033) during exponential growth (Figure 4). In addition, one of two *Klebsiella* isolates (ICBG873, *P*=0.0026) and one of two *Bacillus* isolates (ICBG1751, *P*=0.0008) significantly reduced α-pinene. In addition, other bacterial isolates showed a varying range of reductions of α-pinene between vials, resulting in large variability and lack of significance with the Bonferroni corrected alpha-value. No isolates significantly reduced α-pinene within stationary growth (Figure S2). One of three *Pseudomonas* isolates (ICBG639, *P*<0.0001) significantly reduced β-caryophyllene in the exponential environment. The same *Pseudomonas* isolate (ICBG639, *P*=0.0014) also significantly reduced β-caryophyllene at the stationary phase. Linalool was reduced by two isolates, *Burkholderia* (ICBG637; exponential *P*=0.0036, stationary *P*=0.0008) and *Pseudomonas* (ICBG967; exponential *P*=0.0020, stationary *P*=0.0024). Finally, eucalyptol, the fourth compound tested, was not reduced by any of the isolates tested (Figure S3). In all cases, no breakdown products of the tested PSCs were detected by GC-MS.

**Figure 4.**
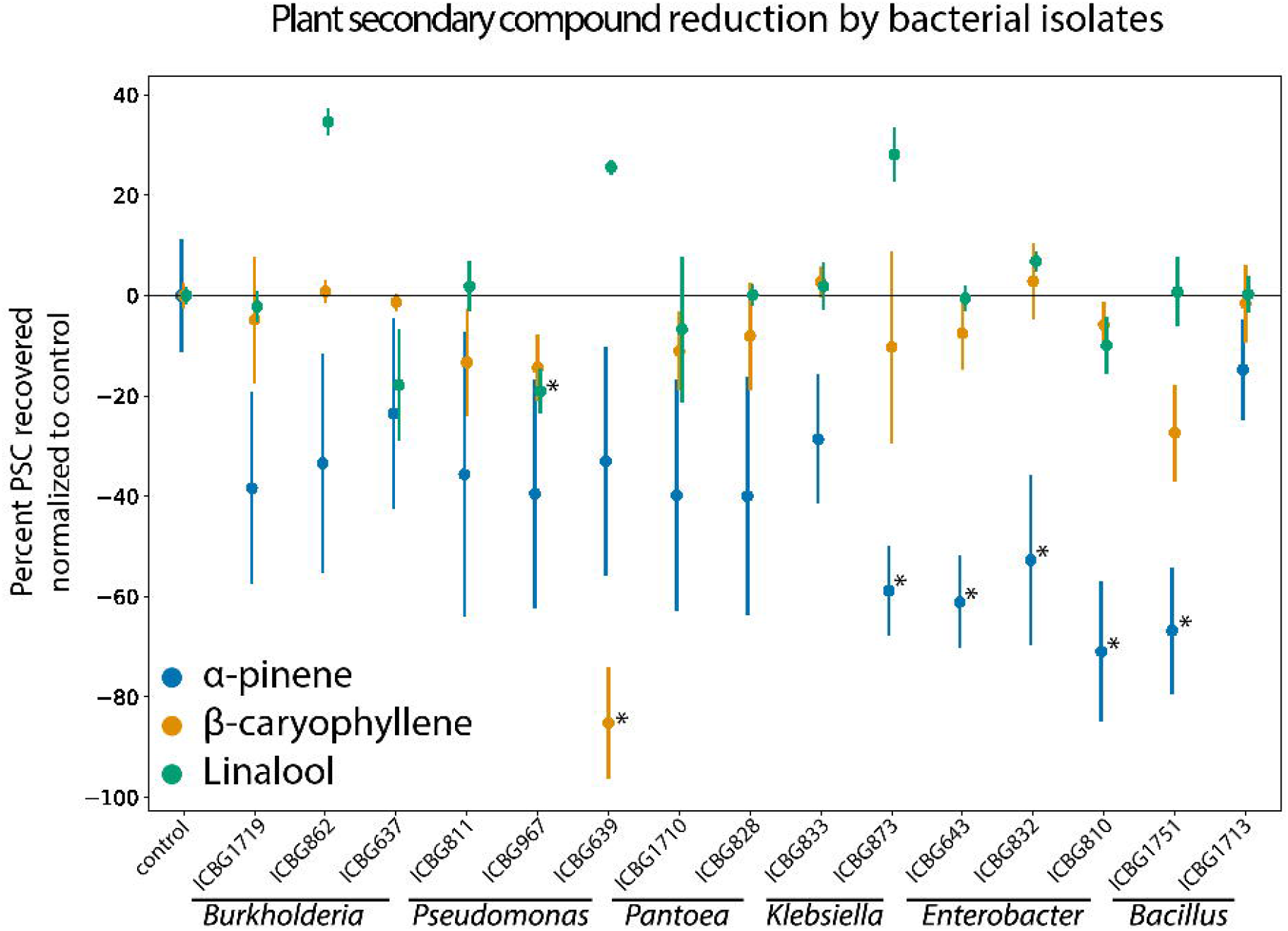
PSC reduction *in vitro* by 15 bacterial isolates. Bacterial isolates are grouped by their genus-level identification on the x-axis. The y-axis is the percent change of PSC recovered compared to a non-bacterial control vial (10% TSB + compound). Each point represents the average of three GC-MS measurements and the bars are the standard deviation of the observations.

##### Headspace sampling of fungus gardens and *L. gongylophorus* with PSC

When sub-colonies from three *A. cephalotes* colonies were exposed to α-pinene (Figure 5A), there was significant reduction of α-pinene in the fungus garden samples at 12 hours (only α-pinene vs fungus garden + α-pinene, *P*<0.0001; cotton + α-pinene vs fungus garden + α-pinene, *P*=0.1191), 24 hours (only α-pinene vs fungus garden + α-pinene, *P*<0.0001; cotton + α-pinene vs fungus garden + α-pinene, *P*<0.0001) and 36 hours (only α-pinene vs fungus garden + α-pinene, *P*<0.0001; cotton + α-pinene vs fungus garden + α-pinene, *P*<0.0001), compared to most control vials (mixed regression model with time and treatment as fixed effects and ant colony as random effects, α = 0.05). In addition, the 36-hour sub-colonies had significantly reduced α-pinene compared to the 12-hour sub-colonies (*P*<0.0001) and the 24-hour sub-colonies (*P*<0.0001). *L. gongylophorus* strains were tested in a similar fashion. Vials containing *L. gongylophorus* were exposed to α-pinene and the headspace was measured after 36 hours of exposure (Figure 5B). Compared to the control vials, *L. gongylophorus* strains did not reduce α-pinene significantly (*P*=0.2786, Welch two sample *t*-test).

**Figure 5.**
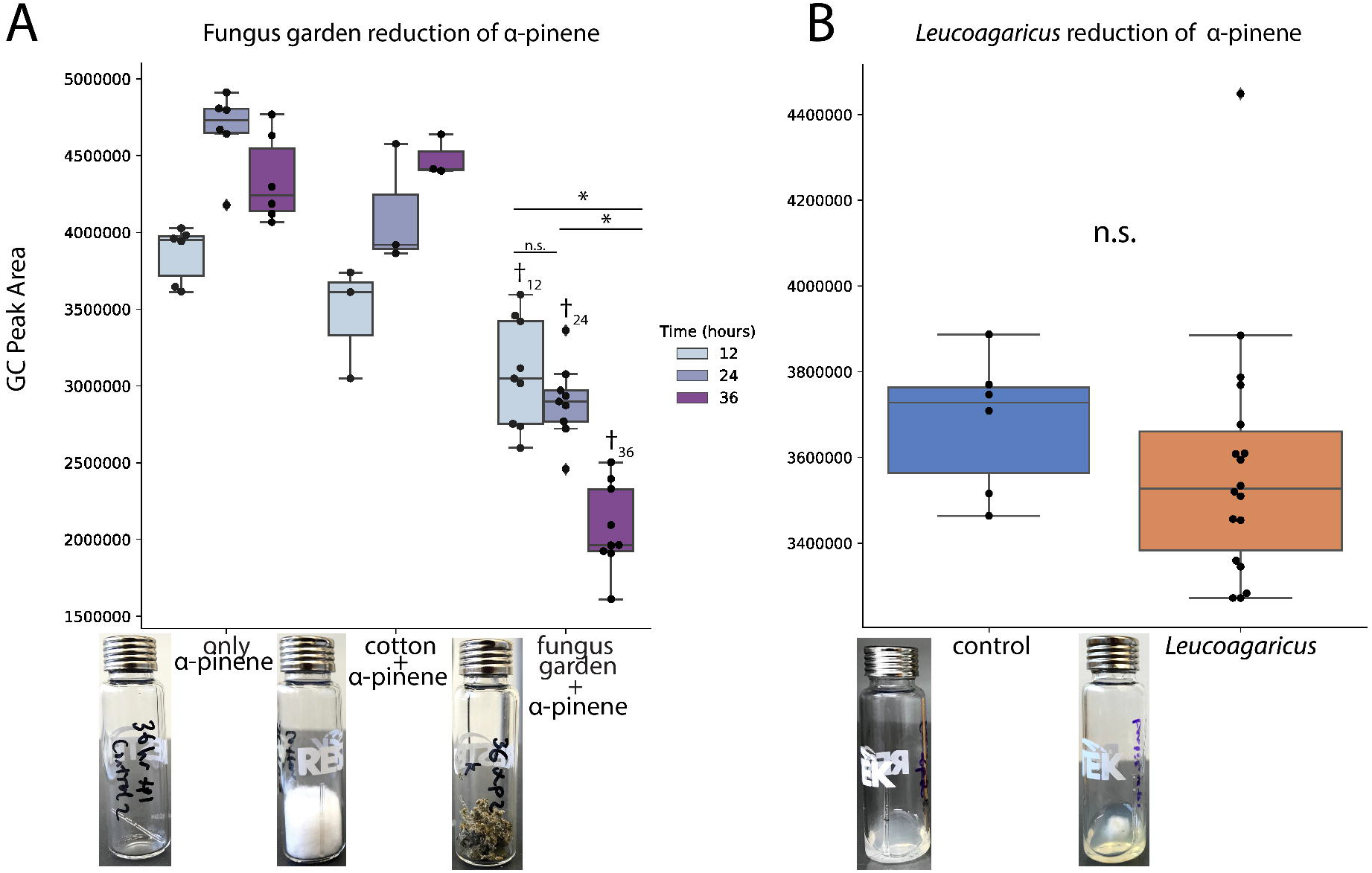
Measurement of α-pinene in the headspace of fungus gardens and *L. gongylophorus* over time. We exposed three sub-colonies of three *Atta cephalotes* colonies to α-pinene for 12, 24, 36 hours and measured reduction of α-pinene levels in the headspace (A). We grew *L. gongylophorus* from the lab colonies, as well as the *L. gongylophorus* strains tested in Figure 2, on PDA and then exposed the strains to α-pinene for 36 hours using micropets flame-sealed at one end. Headspace measurements were collected and compared to a vial with only PDA (B). In (A) and (B) each point represents a single measurement and is superimposed on top of a boxplot. The y-axis indicates the area under the curve with a retention time of ~3.2 minutes, indicating the level of α-pinene in the headspace of each vial. The asterisks (*) indicate significance (*P*< 0.05) between the fungus garden samples at different time points. The cross symbol (†) indicates significance (*P*< 0.05) between the fungus garden sample and the control vials at the same time point. The pictures are examples of the experimental set up for each treatment.

When sub-colonies from five *A. cephalotes* colonies were exposed to linalool (Figure S5A), there was a significant reduction of linalool in the headspace between controls and 12-hour sub-colonies (only linalool vs fungus garden + linalool, *P*<0.0001; cotton + linalool vs fungus garden + linalool, *P*=0.0728), 24-hour sub-colonies (only linalool vs fungus garden + linalool, *P*<0.0001; cotton + linalool vs fungus garden + linalool, *P*=0.0021), and 36-hour sub-colonies (only linalool vs fungus garden + linalool, *P*<0.0001; cotton + linalool vs fungus garden + linalool, *P*=0.0049). We do not see any significant differences between the 12-hour, 24-hour, and 36-hour linalool headspace levels. For the headspace sampling with *L. gongylophorus*, linalool was significantly reduced (*P*=0.0036, Welch two sample *t*-test).

##### Metatranscriptomic sequencing of fungus gardens

We detected the expression of α-pinene/limonene degradation encoding genes in the metatranscriptomic dataset (Table S5, Figure S6A). In the first *A. cephalotes* colony (FG1), we detected 33% of the unique genes in the α-pinene/limonene pathway in a range of 0.506-2831 transcripts per million (TPM), and all three shared genes in a range of 0.337-3.051 TPM. Of note, we detected the gene encoding limonene 1,2-monoxygenase (K14733) expression at 0.506 TPM, which is the first step of limonene transformation. In the second *A. cephalotes* colony (FG2), we detected 22% of the unique genes in the α-pinene/limonene pathway with 59.99 TPM and 3724 TPM, and all three shared genes in a range of 0.168-11.54 TPM. In the *A. colombica* colony (FG3), we detected 22% of the unique genes α-pinene/limonene pathway with 9.062 TPM and 2238 TPM, and one of the shared genes with 4.178 TPM. In all three metatranscriptomes monoterpene epsilon-lactone hydrolase (K14731) was expressed, reflecting its presence in the individual bacterial isolate genomes and confirming the gene’s expression *in situ*.

In the metatranscriptomic dataset (Table S5, Figure S6A), we found that FG1 expressed 50% of the unique genes in the geraniol pathway in a range of 0.115-25.47 TPM, including the gene encoding for citronellol dehydrogenase (K13774) at 0.147 TPM, which is involved in the first step of geraniol and citronellol transformation. In FG2, we detected 42% of the unique genes in the geraniol pathway in a range of 0.033-27.18 TPM and in FG3, we detected 16% of the unique genes in the geraniol pathway with 1.334 TPM and 9.208 TPM. We did not detect the expression of any genes involved in the cumate or trans-cinnamate pathways, nor any expression of the 20 cytochrome p450s or *shc*.

## Discussion

Microbes can mediate plant-insect interactions, including influencing a herbivore’s capacity to overcome plant chemical defenses (15). In this study, we show PSC detoxification in leaf-cutter ants as a polymicrobial process with bacterial communities that supplement the degradative capabilities of the ants’ fungal mutualist, *L. gongylophorus*. Building on previous literature (38–42), we demonstrate that ant-associated strains of *L. gongylophorus* and *Leucoagaricus* sp. have variable ability to tolerate and degrade PSCs (Figure 2, Figure 5), suggesting the necessity for additional symbionts to detoxify the diverse defensive compounds that leaf-cutter ants will encounter as generalist herbivores. Bacteria we commonly isolate from the fungus gardens of leaf-cutter ants both tolerate and degrade PSCs *in vitro* (Figure 3 - 4). Whole genome sequencing of these bacteria revealed the presence of genes involved in PSC degradation (Figure 3), which is further supported by our analysis of existing fungal garden metagenomes and metatranscriptomes (Figure S6, Table S4). Through ant sub-colony experiments, using garden substrates with and without PSC added to the environment, we found *in vivo* degradation of PSC by fungus gardens (Figure 5). Taken together, our findings indicate that the symbiotic microbes within the fungus gardens of leaf-cutter ants can detoxify plant defensive compounds, which may contribute to the overall success of these generalist herbivores.

We show that strains of ant cultivated *L. gongylophorus* and *Leucoagaricus* sp. vary in their sensitivity to and degradative ability towards different PSCs. For example, in tolerance assays, linalool completely inhibited the growth of almost all *L. gongylophorus* and *Leucoagaricus* sp. strains, while most strains grew unperturbed in the presence of farnesol (Figure 2). These results fit with the findings of other studies investigating the toxicity of various PSCs to the fungal cultivars of leaf-cutter ants (38–41). Although *L. gongylophorus* appear sensitive to many PSCs in Petri plate assays, the biosynthetic capacity to degrade some PSCs are present in their fungal genomes, including phenols through production of laccases (42). In addition to our tolerance assays, we also measured the ability of *L. gongylophorus* to degrade α-pinene or linalool by measuring the headspace of vials containing the fungal mutualist and PSC. In these experiments, *L. gongylophorus* did not significantly reduce α-pinene (Figure 5B), but did significantly reduce linalool, which was surprising due to high inhibition by this compound in the plate assay (Figure 2B, Figure S5B). The observed degradation of linalool by *L. gongylophorus* may be due to a difference in dosage, as the tolerance assays contained high concentrations of compound, whereas the headspace sampling had lower amounts of compound and *L. gongylophorus* obtained a higher biomass (grown for longer on agar plates). Overall, the variation of *L. gongylophorus* to tolerate and degrade PSC implies that the bacterial community is potentially involved in reducing PSCs that would otherwise inhibit the fungal mutualist.

Fungus garden bacteria demonstrated high tolerance of PSC and significant degradation of α-pinene, β-caryophyllene or linalool. Specifically, we tested bacterial isolates for *in vitro* PSC tolerance and/or degradation, including strains representing the dominant fungus garden microbiome: *Burkholderia, Enterobacter, Klebsiella, Pantoea,* and *Pseudomonas* (26–28, 31). In the tolerance assay, we saw widespread resistance to the PSCs by most bacterial genera, even in the cases of isolates not predicted to contain degradation genes of that PSC’s pathway (Figure 3A-B). For example, all isolates except for *Acinetobacter* and *Bacillus* were completely resistant to *p*-cymene, perhaps due to an alternative to degradation, such as efflux pumps (43). In addition, while most isolates, except for *Bacillus*, tolerated the eight PSCs, we saw marked difference in the isolates ability to degrade PSCs. One isolate of *Bacillus*, while inhibited in our tolerance plate assay, significantly reduced α-pinene in our GC-MS experiment (Figure 4). This could be due to a difference in dosage between the two experiments, which has been shown to have an effect on bacterial tolerance and degradation of PSCs (9). *Burkholderia* and *Pseudomonas,* which both significantly reduced PSCs in our GC-MS assay, have been implicated in PSC degradation in other systems, including bark and mountain pine beetles (8, 44). While not cultured in our study, bacteria in the genera *Serratia* and *Rahnella* isolated from bark beetles were found to degrade PSCs. *Serratia* and *Rahnella* have been detected in leaf-cutter ant fungus garden sequencing studies (32, 33, 36, Figure S4), suggesting that other isolates within fungus garden bacterial communities may be involved in degrading PSC. Finally, no breakdown products were detected via GC-MS, which could be a result of complete degradation of the compounds into components of central metabolism (45, 46).

Garden bacterial genomes and fungus garden metagenomes and metatranscriptomes indicate that genes involved in PSC degradation are present and expressed within the fungus garden bacterial community. Specifically, the presence of cytochrome p450 encoding genes, cumate, trans-cinnamate, and *p*-cymene degradation pathways, as well as the presence and *in situ* expression of gesnes in the α-pinene/limonene and geraniol degradation pathways indicate that garden bacteria are predicted to metabolize PSC. In addition, the presence of the gene encoding *shc*, which synthesizes hopanoids and increases membrane stability (47), in *Asaia* and *Burkholderia* isolates could explain the ability to tolerate stressful conditions, such as growth in the presence of PSC. With the available data, we were able to predict that both individual garden bacteria isolates and garden bacteria metagenomes contained the genes necessary to degrade or transform PSC that could harm the fungus gardens. Of note, the higher completeness of pathways observed in the metagenomes suggests that while individual strains may not have the entire pathway for the degradation of a particular compound, as a community, garden bacteria have the capabilities to reduce PSC within the fungus garden. The concept of facilitation (48), where one organism will accidentally or purposefully benefit from another, in microbial communities has been widely explored, including in the presence of toxins (49). Facilitation between microbes has been observed in other insect systems, such as the gut microbiomes of honey bees, which contain microbial symbionts that have complementary abilities (50) and cross-feeding between microbial community members has been observed (51). Further experiments are necessary to address the microbial community potential to complement each other in the degradation of various plant secondary compounds and if different community compositions would impact plant intake by the fungus gardens.

We further assess the contribution of microbes within ant gardens to degrade PSC by exposing pieces of *A. cephalotes* fungus gardens to α-pinene or linalool and measuring the headspace over time. In the presence of both α-pinene and linalool, the fungus gardens significantly reduced the amount of PSC in the headspace compared to vials with no fungus garden. After 12 hours of exposure, the headspace of fungus gardens contained α-pinene and linalool levels 20% and 52% lower, respectively, than vials containing only PSC (i.e., no fungus garden). α-pinene levels in the headspace of vials containing fungus garden decrease over time, with levels at 36 hours post-exposure being 28% lower than the 24-hour timepoint. In contrast, linalool levels in the vials containing fungus garden remain stable after 12 hours, suggesting there may be a limit to the degradation possible with this compound. Finding significant *in vivo* reduction of PSC by microbes in the fungus garden, in combination with the *in silico* and *in vitro* evidence of fungal and bacterial PSC tolerance and degradation, indicates the ability of the leaf-cutter ant external digestive system to mitigate the presence of PSC.

Like other herbivorous insect systems, the gut microbiome of leaf-cutter ants is demonstrably important for dictating palatable plant substrate. Through a combination of *in vitro* and *in vivo* approaches, our study provides evidence that the consistent bacterial community in fungus gardens contributes to the detoxification of PSCs, potentially enabling leaf-cutter ants to forage from a wide variety of plant sources. Overall, the symbioses formed between fungal and bacterial symbionts with leaf-cutter ants demonstrates the intricacy and nuance with which microbes serve as an interface between herbivores and the plants they consume and contributes to the ecological success of these systems.

## Materials and Methods

### *L. gongylophorus* and *Leucoagaricus* sp. tolerance of PSC

We selected compounds for testing based on leaf extracts from plant families that have been foraged by leaf-cutter ants (17), detection of terpenes in fungus gardens of *A. laevigata* (31), and commercial availability: 98% (1R)-(+)-α-pinene (Acros Organics), >90% β-caryophyllene (TCI), 99% eucalyptol (Sigma-Aldrich), 95% farnesol (Sigma-Aldrich), 96%(S)-(-)-limonene (Sigma-Aldrich), 97% linalool (48.2% (R)-(−)-linalool/51.8% (S)-(+)-linalool) (Sigma-Aldrich), 99+% *p*-cymene (Acros Organics), and 85% terpinolene (Sigma-Aldrich). The eight PSC were tested against five strains of fungal cultivar from *A. sexdens*, *A. laevigata*, *A. bisphaerica*, *A. capiguara*, and *P. diversus* colonies (isolation information in Table S1). Of note, while *P. diversus* is not within the leaf-cutter ant lineage and largely collects substrate like seeds, insect frass, and dry plant debris for its garden, the species has been observed occasionally collecting fresh leaf and flower material as substrate for its fungus garden (52, 53). We put a 6 mm fungal plug of *L. gongylophorus* or *Leucoagaricus* sp. onto a 60 mm Oxoid Malt Extract Agar (OMEA; per L: 30 g malt extract, 5 g mycological peptone, 15 g agar) plate and cultured for two weeks at ambient temperature. Then, for each PSC, we placed a sterile disc with 15 μL of undiluted compound that had been allowed to dry in a biological safety hood for five min 1 cm from the edges of fungal growth. We tested each compound in triplicate (three plates per PSC per cultivar) and inhibition was monitored over the course of two weeks, with pictures being taken on day 2 and 14. Inhibition was determined by a qualitative scale where 0 represents no inhibition, 1 represents no/slight inhibition at day 2 and normal growth by day 14 (compared to control), 2 represents no/slight inhibition at day 2 and resume slow growth by day 14 (compared to control), 3 represents mostly inhibited by day 2, no additional growth by day 14, 4 represents complete inhibition at day 2 and day 14 (Figure 2A). Inhibition was measured by the same individual based on direct observation.

### Sampling and bacterial isolations

We collected fungus-farming ant colonies in January 2017 in the following general locations: Anavilhanas, AM; Ducke Reserve, AM; Itatiaia, RJ; Botucatu, São Paulo State; Ribeirão Preto, São Paulo State. Details regarding the exact GPS coordinates and environment of the samples can be found in Table S1. In the field and lab setting, we vortexed pieces of fungus garden from the middle in 1X PBS, which we then pipetted on to Yeast Malt Extract Agar (YMEA; per L: 4 g yeast extract, 10 g malt extract, 4 g dextrose, 15 g agar). Garden bacteria were isolated from multiple fungus gardens of the two genera of leaf-cutter ants, *Atta* sp. and *Acromyrmex* sp., as well as three genera of other fungus-growing ants, *Paratrachymyrmex* sp., *Cyphomyrmex* sp., and *Apterostigma* sp. (Table S1). We obtained pure isolates after several rounds of subculturing based on morphology for a total of 317 isolates. We identified 117 isolates to genus-level by 16S *rRNA* gene sequencing as previously described (54). Briefly, we lysed colonies and PCR was performed with 16S rRNA primers 27F (5’-GAGAGTTTGATCCTGGCTCAG-3’) and 1492R (5’-GGTTACCTTGTTACGACTT-3’). We sequenced samples using Sanger sequencing at the University of Wisconsin – Madison Biotech Center (Madison, WI) and analyzed using 4Peaks and CLC Sequence Viewer 7. We matched the 16S *rRNA* gene sequence using BLAST (55) and the SILVA database(https://www.arb-silva.de) for the nearest genus-level identification.

### DNA Extraction, Assembly, and Annotation

We selected forty-two bacterial isolates for whole genome sequencing based on belonging to genera known to be abundant and consistent in fungus gardens (*Burkholderia, Enterobacter, Klebsiella, Pantoea, Pseudomonas*) or belonging to genera less common in the fungus garden (*Acinetobacter, Asaia, Bacillus, Chitinophaga, Chryseobacterium, Comamonas*). We extracted DNA from 42 bacterial isolates using the Promega Wizard Genomic DNA Purification Kit using the Gram-Negative and Gram-Positive Bacteria Protocol. We used Qubit BR dsDNA kit (Invitrogen, USA) for quality control measures. We prepared genomic DNA libraries for Illumina MiSeq 2×300bp paired-end sequencing by the University of Wisconsin-Madison Biotechnology Center. We corrected reads with MUSKETv1.1 (56), merged paired-ends with FLASH v1.2.7 (57), and assembled with SPAdes 3.11.0 (58). Genome statistics can be found in Table S2. We determined species-level identification by uploading the genomes to JSpeciesWS (59), performing a Tetra Correlation Search, and taking the first result. If there were conflicts in the top 5 results (i.e., different genera), the top 5 genomes were pulled and ANI with pyani (60) using the ANIm analysis was performed. Then, we selected the genome with the highest percent similarity as our isolate’s taxonomic status. We identified some isolates differently based on whole genomes from the 16S taxonomic classification, such as one *Burkholderia* isolate (ICBG641) belonged to the genus *Paraburkholderia* (Table S2).

We identified predicted proteins putatively involved in PSC degradation from each genome in one of two ways: 1) Using the KEGG Automatic Annotation Server (KAAS). Specifically, we investigated genes encoding enzymes putatively involved in monoterpene degradation or aromatic compound degradation, as defined in the KEGG limonene and α-pinene degradation pathway (ko00903), geraniol degradation (ko00281), degradation of aromatic compounds (ko01220, modules 00419,00539, 00545) after annotation. 2) DIAMOND v0.9.21.122 (61) BLASTP against the Uniprot Swiss-Prot and TrEMBL databases (www.uniprot.org/downloads), downloaded on July 18, 2019. We only kept the top 5% of hits (--top 5) that had an e-value below 1e-05 for each query sequence. Then, using the grep command, we looked for the following Uniprot accession numbers that corresponded to 20 cytochrome p450s (36): P00183, P14779, Q2L6S8, P18326, Q59079, Q59831, Q06069, P53554, P33006, U5U1Z3, A9F9S4, Q59723, Q59990, Q9K498, Q53W59, Q8VQF6, A9FZ85, Q88LH7, Q88LH5, Q88LI2, Q65A64; the 20 genes in the diterpene degradation cluster (62): Q9X4W9, Q9X4W8, Q9X4X8, Q9X4X7, Q9X4X6, Q9X4X5, Q9X4X4, Q9X4X2, Q9X4X1, Q9X4X0, Q9X4W7, Q9X4W6, Q7BRJ3, Q7BRJ4, Q7BRJ5, Q7BRJ6, Q7BRJ7, Q7BRJ8, Q7BRJ9, Q9X4X; saxA (63): A0A0N7FW12; squalene-hopene cyclase (47): P33990, P54924, P33247. We also performed DIAMOND BLASTP using a custom database with solely these gene sequences with a query coverage cut-off of 75% (--query-cover 75) and e-value cut-off of 1E-05. If there was alignment in both the Uniprot analysis and the custom analysis, then genes were predicted to be present. The annotation methods were used on individual bacterial genomes, one *L. gongylophorus* genome (BioProject: PRJNA179280), one *Leucoagaricus* sp. genome (BioProject: PRJNA295288), and publicly available leaf-cutter ant garden bacteria metagenomes from Brazil (Gold Analysis Project ID: Ga0157357 – Ga0157368). For the metagenomes, we used the metagenomes KAAS option instead of the complete/draft genome KAAS option.

### Bacterial tolerance of PSC

We tested the effect of the eight PSC on 46 bacterial isolates using Whatman 6 mm discs. Bacterial isolates were grown overnight (16-24 hours) until turbid (OD_600_ = ~1-2). We spread 100 μL of overnight culture on YMEA plates using a glass cell spreader. We deposited a disc with 15 μL of PSC in the center of the bacterial lawn. Each PSC was done in triplicate (3 plates per plate secondary compound per bacterial isolate). After 48 hours, we took pictures of the plates using an Epson scanner and then uploaded the photos into Fiji (64). We used Fiji Version 1.0 to measure the zones of inhibition caused by each PSC (in centimeters). We calculated the average of the three zones of inhibition and then scaled all the zones of inhibition in reference to the largest zone observed so that 0 indicates inhibition (zone of inhibition = 3 cm) and 1 indicates no inhibition (100% growth; zone of inhibition = 0 cm)

### Gas Chromatography-Mass Spectrometry (GC-MS) of bacterial isolates incubated with PSC

We prepared fifteen bacterial isolates representing highly resistant genera (*Burkholderia, Enterobacter, Klebsiella, Pantoea, Pseudomonas*) and inhibited genera (*Bacillus*) two ways for GC-MS: addition of compound during exponential growth or stationary growth. For both methods, bacterial isolates were grown overnight (16-24 hours) in 10% tryptic soy broth (TSB). All shaking was done at 300 rpm at room temperature. All experiments included an extra vial to read the OD_600_ to ensure bacterial growth in the presence of compound, which also served to confirm earlier patterns of compound tolerance by bacterial isolates (data not shown). *(A) Exponential phase* We diluted overnight cultures to OD_600_ = 0.08. We inoculated the appropriate amount of overnight culture into vials containing 10% TSB and 2.5 μL/mL of one of four PSC, α-pinene, β-caryophyllene, eucalyptol, linalool, that were added using a glass manual GC syringe (10 μL, Thermo Scientific). We left the bacterial cells and plant compound shaking for another two days at room temperature. This was done with 15 bacterial isolates representing six genera, as well as negative controls (no bacteria), in triplicate (16 × 4 compounds × 3 replicates). *(B) Stationary phase* We pipetted 10 μL of overnight culture into 987.5 μL of 10% TSB in vials. After two days of incubation at room temperature while shaking at 300 rpm, we added 2.5 μL/mL of one of four PSC (α-pinene, β-caryophyllene, eucalyptol, linalool) directly to the vials using a glass manual GC syringe (10 μL, Thermo Scientific). Then, we left the bacterial cells and plant compound shaking for another two days at room temperature. This was done with the same number of samples listed above.

For both method A and B, we extracted PSC by pipetting 1 mL of hexane into each vial and shaking the vials overnight. We removed 500 μL of the hexane-PSC phase and put into new vials containing 500 μL hexane and 5 μL/mL of the internal standard toluene. We then analyzed the abundance of each PSC using GC-MS. Specifically, the GC system consisted of a Thermo Fisher Trace 1310 Gas Chromatograph coupled with Thermo ISQ LT Single Quadrupole Spectrometer. We injected 1 μL of each mono-/sesqui-terpene sample directly, with a split flow ratio of 30:1. We used an oven profile of 40 °C, followed by a ramp of 3 °C min^−1^ to 115 °C (monoterpenes) or 130°C (sesquiterpenes) and then 30 °C min^−1^ to 250 °C with a 2 min hold. We integrated and analyzed peaks using the Chromeoleon Chromatography Data System Software.

We integrated and standardized signal peaks from the GC based on the internal standard toluene (peak area/internal standard peak area) for each vial. In addition, we used standard curves of the four pure PSC to measure changes in concentration in the samples when compared to controls. We made standard curves to incorporate the possible ranges of concentrations (0 μL/mL to 3.5 μL/mL) within the experiment. We then calculated proportional change of bacterial treatments versus the nonbacterial control. Specifically, we took the average of the non-bacterial control standardized peak areas and subtracted the control average from all the bacteria-compound peak areas. Then, we divided the adjusted value by the non-bacterial control average to obtain the percent change [(bacterial standardized peak area – average of control standardized peak areas)/average of control standardized peak area]. We then analyzed the standardized values in JMP Pro 13 by performing one-sampled Student’s *t*-tests for each compound with a null hypothesis of μ=0, representing no change between compound abundance in bacteria-treated and the non-bacterial control. Since we were performing 15 separate statistical tests for each compound (between non-bacterial control and each of the 15 bacterial isolates), we used a Bonferroni correction to avoid false positives (α=0.05/15=0.0033).

### Headspace sampling of fungus gardens with PSC

*Atta cephalotes* colonies (Table S3) that have been maintained in lab from one to seven years were used in this experiment to create sub-colonies. 16S *rRNA* gene amplicon sequencing was used to confirm that the bacterial genera in these fungus gardens were consistent with isolates used throughout this study (Text S1, Figure S4B). We prepared twenty mL 18 mm Restek (Bellefonte, PA, USA) vials (cat#23082) with magnetic screw-thread caps (cat#23090) three ways with α-pinene or linalool: (1) Empty vials with a 20 μL Accu-Fill 90 micropet cut to 2.54 cm and flame-sealed at one end containing 1 μL of PSC (n=6). (2) Vials with approximately 0.3-0.4g of cotton with a 2.54 cm micropet flame-sealed at one end containing 1 μL of PSC (n=3). (3) Vials with approximately 0.3g of fungus garden material with all ants manually removed and a 2.54 cm micropet flame-sealed at one end containing 1 μL of PSC (n=3 subsamples × 3-5 different *A. cephalotes* colonies). We used material from three colonies in the α-pinene experiment and five colonies in the linalool experiment. We also prepared samples of vials with only fungus garden (i.e. no exposure to PSC) during certain runs to ensure that there were no detectable PSC innate to the system. This was done for three separate time points based on exposure to a PSC: 12 hours post-exposure, 24 hours post-exposure, and 36 hours post-exposure. At these given time points, we destructively sampled each respective set of vials with a Shimadzu HS20 Headspace Sampler coupled to a Shimadzu GC-2010 Plus with a flame ionization detector. Specifically, we loaded vials into the headspace sampler and injected into a column with a 50:1 split flow ratio. For the vials with α-pinene, the headspace sampler and oven were at 60 °C, followed by a 20 °C/min ramp up to 140 °C. For the vials with linalool, which has a higher boiling point than α-pinene, the headspace sampler and oven were kept at 70 °C, followed by a 25 °C/min ramp up to 205 °C. Then, we identified compounds using retention time (α-pinene=3.2 minutes, linalool=6.2 minutes) and calculated areas under the curve in Shimadzu’s LabSolutions software to determine the relative difference in α-pinene or linalool between vials. We chose α-pinene and linalool are compounds that the garden bacteria can degrade and are inhibitory against *L. gongylophorus* and *Leucoagaricus* sp.

Since we took subsamples (sub-colonies) from each *A. cephalotes* colony (3 subsamples × 3 time points × 5 colonies), we employed a linear mixed-effects model to account for the correlation (non-independence) between subsamples. Specifically, to test if ant colony had an effect on the observed value, we used the lmer package v. 3.1-0, holding time and treatment as the fixed effects and ant colony as the random effect. Before the analysis, we divided the values by 1,000,000 to rescale the response for the lmer optimization procedure. For the α-pinene treatment, the colony variance is reported as 0, indicating that the variability with respect to ant colony is much smaller than the variability with respect to the residual error. For the linalool treatment, the colony variance was 0.000226, indicating that some of the variability observed was due to the sampling from different colonies. We then used the estimated marginal means (EMMs) with the emmeans package v. 1.3.5 for linear regression analysis of the data, using the pairs() method. Marginal means were compared pairwise between exhaustive two-way level combinations of treatment (control, cotton, fungus garden) and of time (12hr, 24hr, 36hr). Assumptions of normality, linearity, and homoscedasticity for linear regression were examined by plot diagnostics and were met for each analysis. All the code used in this analysis is available at github.com/cfrancoeur/PSC.

### Headspace sampling of *L. gongylophorus*

We isolated *L. gongylophorus* strains by plating small pieces of healthy fungus garden from laboratory colonies on Potato-Dextrose Agar (PDA). Laboratory fungus-farming ant colonies are kept in a temperature-controlled 28 °C room in separate large plastic containers. We used five *A. cephalotes* colonies collected over the course of several years (2012-2018) from Costa Rica (Table S3). In addition, several isolates from Brazilian *Atta* gardens used in the fungal cultivar tolerance experiment (Figure 2) were included.

We pipetted 2 mL of PDA into 20 mL 18 mm Restek vials with magnetic screw-thread caps and left to solidify on a slant. Then, 3×3 mm pieces of freshly growing *L. gongylophorus* strains were placed onto the slant and grown for one month at room temperature in the dark. We prepared three vials for each of the *A. cephalotes* cultivars (n=5 strains × 3 vials × 2 compounds) and we prepared one vial for three additional *L. gongylophorus* strains: AB1, AL2, AS1 (n=3 strains × 1 vial × 2 compounds). After the month of growth, we filled 20 μL Accu-Fill 90 micropets (Becton, Dickinson and Company, N.J.) cut to 2.54 cm and flame-sealed at one end with (A) nothing (B) 1 μL α-pinene or (C) 1 μL linalool. We then put the filled micropets into the vials and after 36 hours of exposure, we analyzed the headspace of the vials with the same methodology described for the sub-colony headspace sampling. We statistically compared signal peaks using a Welch two sample *t*-test, comparing the peaks from the control vials to the vials containing *Leucoagaricus* sp.

### Metatranscriptomic sequencing of fungus gardens

We collected samples directly from the field into RNAlater buffer. We took samples from the top sections of three different colonies: two *A. cephalotes* colonies from La Selva Biological Station, Costa Rica and one *A. colombica* colony from Golfito, Costa Rica (Table S4). Total RNA extraction was identical to a method previously described (65). We performed cDNA library construction and Illumina HiSeq2000 sequencing at the University of Wisconsin Biotechnology Center (Madison, WI). We uploaded the metatranscriptomes to MG-RAST and processed the reads with their SOP (66). We downloaded the reads post-processing (quality reads) and then analyzed metatranscriptomes using prodigal V2.6.2 (67), DIAMOND v0.9.21.122, and kallisto v. 0.43.1 (68). First, we ran prodigal on the assembled nucleotide files of 12 garden bacteria metagenomes from Brazil (downloaded from JGI, accession numbers provided in annotation methods above) with the metagenomic flag (-p meta). Then, we created a kallisto index with all of the combined prodigal garden bacteria metagenome output. We used the kallisto quant command to pseudo-align the garden bacteria index against the metatranscriptome reads. This gave a transcripts per million (TPM) value of bacterial transcripts in the metatranscriptome. Then, we used DIAMOND to blastp the metagenome coding regions against the Uniprot and KEGG databases described above. Using grep, we found the genes of interest (same as in the bacteria isolate annotation) and connected the gene of interest, the metagenomic transcript it mapped to, and the TPM in the metatranscriptome. For genes with multiple transcripts and different TPMs, we recorded the unique values (Figure S6) and summed the TPMs for total expression (Table S5). We also did the same workflow for four housekeeping genes (*gyrB*: K02470, *rpoB*: K03043, *rpoD/sigA*: K02086, *rpsL*: K02950).

## Data Availability

All sequencing data has been uploaded to NCBI under the following BioProject numbers: PRJNA564151, PRNJNA429666, PRNJNA429667, PRNJNA429668, PRNJNA565936, PRJNA577467. Individual accession numbers for each dataset are included in Table S2, Table S3, and Table S4.

---

**Figure S1.** Plant secondary compound tolerance of bacteria isolated from fungus gardens of fungus-farming ants. The dataset includes the 31 bacterial isolates from Figure 3 in the main text, with 15 additional isolates that were tested for tolerance but do not have whole genome sequences. Blue indicates resistance to the eight compounds tested and red indicates high inhibition.

**Figure S2.** Plant secondary compound degradation by bacterial isolates exposed to compounds after growing for 48 hours (stationary phase).

**Figure S3.** Plant secondary compound degradation by bacterial isolates exposed to eucalyptol. These data are the results from eucalyptol being added to diluted bacteria (exponential), as well as from eucalyptol being added after bacteria had been growing for 48 hours (stationary). None of the isolates could reduce eucalyptol during either condition.

**Figure S4.** Fungus garden sub-colonies from five *A. cephalotes* colonies were exposed to 4 conditions: no treatment, addition of α-pinene (2 doses), addition of linalool (2 doses), and removal of ants (total of seven samples per colony). 16S rRNA amplicon sequencing of the fungus gardens from each colony revealed no detectable changes in the bacterial communities between treatments. However, taxa from laboratory colonies are consistent with previous metagenomic and culturing studies done with fungus gardens collected directly in the field both at the phylum level (A) and genus level (B).

**Figure S5.** We exposed five sub-colonies of *A. cephalotes* to linalool for 12, 24, and 36 hours and then sampled the headspace at each time point. Fungus garden vials had significantly reduced amounts of linalool compared to empty vials and vials with cotton (A). When *L. gongylophorus* was grown is isolation and exposed to linalool for 36 hours, there was a significant decrease in linalool compared to the control vials (B).

**Figure S6.** Gene expression of plant secondary compound degradation genes (A) and housekeeping genes (B). The source of transcripts came from a garden bacteria metagenomic dataset and therefore different species will be expressing genes at different levels. The variation of transcript expression is displayed as scatterplots, color-coded by metatranscriptome sample. Any gene that had at least 10 transcripts being expressed has a violin plot super-imposed onto the scatterplot. See Table S5 for the sums of all the individual expression values.

**Table S1.** Collection information for fungal isolates used in tolerance assay and bacterial isolates chosen for whole-genome sequencing.

**Table S2.** Genome statistics and accession numbers for 42 bacterial isolates.

**Table S3.** Laboratory colony collection information used for headspace sampling and 16S rRNA amplicon experiments.

**Table S4.** Fungus garden collection information for metatranscriptomic sequencing.

**Table S5.** Sum of bacterial gene expression of plant secondary compound degradation and housekeeping genes. See Figure S6 for individual TPM for different bacterial transcripts.

**Text S1.** Supplemental methods, results, and discussion for 16S rRNA amplicon sequencing and dosage experiment (bacterial community changes when fungus gardens are exposed to PSC).

## Acknowledgments

We are grateful to members of the Landick Lab and Amador-Noguez Lab at UW-Madison, Sherry Cao and Mehmet Tatli, for training on the Shimadzu Headspace sampler. We thank Michael Liou and Michael Howe for their assistance with the statistical analyses. We thank Ted Schultz for species-level identification of our *P. diversus* fungus-farming ant specimen. We thank Reed Stubbendieck and Margaret Thairu for constructive comments on the manuscript. We also thank the staff and scientists at the La Selva Biological Station for allowing field collections of *A. cephalotes* and the scientists Allan Artavia and Miguel Pacheco in the Pinto-Tomás lab at University of Costa Rica for their assistance with field collections. We are grateful to the Comisión Institucional de Biodiversidad of the University of Costa Rica (Resolución N° 009) and the Ministerio de Ambiente y Energía for providing collection permits in Costa Rica. Finally, we thank Heidi Horn for training on collecting fungus-farming ant colonies. This material is based upon work supported in part by the Great Lakes Bioenergy Research Center, U.S. Department of Energy, Office of Science, Office of Biological and Environmental Research under Award Numbers DE-SC0018409 and DE-FC02-07ER64494, the National Institutes of Health grant U19 TW009872, the National Science Foundation grant DEB-1927155, and Vicerrectoría de Investigación (810-B0-501), Universidad de Costa Rica, Ministerio de Ciencia, Tecnología y Telecomunicaciones (MICITT), Costa Rican Government and Florida Ice and Farm Company.

## Author contributions

C.F., L.K., K.G., A.J.B, R.D.M-S., A.A.P-T., K.K-R., C.R.C designed research; C.F., A.J.B., R.D.M-S. performed research; C.F., K.G. analyzed data, and C.F., L.K., K.G., C.R.C. wrote the paper.

