## Supplemental Text 1 for "Bacteria contribute to plant secondary compound degradation in a generalist herbivore system"

Supplemental Methods and Results/Discussion for “Bacteria contribute to plant secondary compound degradation in a generalist herbivore system”

**Materials and Methods**

**Bacterial community changes when fungus gardens are exposed to PSC*.*** Two different doses of compound were used in this experiment: 5 µL in 5.08 cm 50 µL capillaries with one end flame-sealed and 25 µL in 5.08 cm 100 µL capillaries with one end flame-sealed. For each dose there were three treatment groups: (A) control (no compound) (B) α-pinene and (C) linalool. For each treatment, approximately 5 g of the top-middle layer of fungus garden were taken from the same five *Atta cephalotes* colonies used in the headspace experiment (5 colonies x 3 treatments x 2 doses = 30 sub-colonies). The fungus garden piece was placed in a 60 mm glass petri dish to ensure stability and to make final collection easier. Then, the fungus garden piece was placed into a 12 oz glass jar with metal lids (Nakpunar, New Jersey, USA). When all fungus garden pieces had been randomly assigned a treatment for each dose, glass capillaries containing either (A) nothing (B) α-pinene and (C) linalool were placed in a glass tube to hold the capillary upright, then put in the appropriate jar containing fungus garden.

*DNA extraction + 16S rRNA sequencing* After 48 hours, fungus garden pieces were collected into 50 mL conical tubes and weighed. Total DNA was extracted using a bacterial enrichment method previously described (1). Briefly, the fungus garden material was homogenized using a sterile mortar and pestle. Then, the homogenized material was submerged in 1X PBS containing 0.1% Tween 80, followed by centrifugation for 15 min at 500-700rpm. This results in a layered mixture containing leaf-material and fungal mass at the bottom, and bacteria in the middle/top. This washing step was repeated until the bacterial layer became more transparent (about 3-5 times). Then the bacterial sample was spun down, the pellet was resuspended in 1X PBS + .1% Tween 80 and passed through a 40 µm filter to remove any larger, non-bacterial debris. The filter was flushed with an additional 5 mL 1XPBS + .1% Tween 80 then the entire sample was spun down again. Total DNA was extracted using a Qiagen DNeasy Plant Mini Kit. DNA was submitted to the Biotechnology Center for library preparation and sequencing. Specifically, amplicon libraries spanning the V4 region of the 16S ribosomal gene were constructed and then sequenced using Illumina MiSeq 2x300bp at the University of Wisconsin – Madison Biotechnology Center.

Sequence reads were processed, aligned, and categorized using DADA2 1.12.1 (2). The DADA2 pipeline (<https://benjjneb.github.io/dada2/tutorial.html>) was followed almost exactly in July 2019. The only change was made in the filtering step using altered truncLen and trimLeft parameters since we sequenced 2x300bp reads (truncLen=c(225,280); trimLeft=c(10,10)). The statistical analysis was performed as described in the DADA2 pipeline provided above. Additional microbiome analyses, including the violin plots in Figure S6B, were adapted from <https://bioconductor.org/help/course-materials/2017/BioC2017/Day1/Workshops/Microbiome/MicrobiomeWorkflowII.html>. The adapted code can be found in github.com/cfrancoeur/PSC.

**Data availability.** Sequence data can be found at NCBI under BioProject PRJNA577467.

**Results and Discussion**

**Fungus gardens exposed to α-pinene or linalool did not experience a shift in bacterial community composition.** We exposed sub-colonies of *Atta cephalotes* to a low and high dose of α-pinene or linalool for 48 hours and then extracted bacterial DNA for 16S rRNA amplicon sequencing to determine if there was a shift in bacterial community, as previously observed for mammalian herbivore gut microbiomes (3). At the doses and exposure tested, we did not see a change in abundance of certain expected community members known to degrade PSC (i.e., *Pseudomonas*), compared to the control sub-colonies (data not shown). We chose 48 hours of exposure because a preliminary experiment demonstrated a change in bacterial CFUs in fungus gardens after exposure of PSC for this time (data not shown) and because we saw a decrease of PSC in our headspace experiments (discussed below) within 36 hours. The lack of community changes could be a result of the short exposure time and therefore not enough time for bacterial community turnover. Additionally, we used a large piece of fungus garden to obtain enough bacterial DNA for analysis; however, these degradation events could be happening at localized scales, and therefore the large sample size could have swamped out any local degradation/bacterial growth effects. However, we found another result from this experiment interesting; the bacterial genera observed are the same as genera observed from the fungus gardens of leaf-cutter ants collected and immediately processed in the field (1, 4–6) (Figure S6). The colonies used in the experiments have been in lab for approximately 7 years (RM120223-02), 5 years (CR14) and 1 year (CF180406-01, CF180405-02, HH180403-03), which indicates that fungus garden bacterial members – at the genus-level – are maintained even with a large change in environment: tropical rainforest in Costa Rica to a controlled lab setting in Wisconsin. Specifically, we confirmed that the genera we cultured directly in the field for our *in vitro* assays were found in our laboratory colonies. All isolates tested in the GC-MS experiment that had been cultured from the Brazil colonies were detected in the 16S data except for *Bacillus* (i.e. *Burkholderia, Pantoea, Pseudomonas, Enterobacter,* and *Klebsiella* were present at varying abundances (Figure S6B). This means we can assume that our results from the fungus garden headspace experiments are somewhat relatable to environmental fungus gardens.

1.	Suen G, Scott JJ, Aylward FO, Adams SM, Tringe SG, Pinto-Tomás AA, Foster CE, Pauly M, Weimer PJ, Barry KW, Goodwin LA, Bouffard P, Li L, Osterberger J, Harkins TT, Slater SC, Donohue TJ, Currie CR. 2010. An insect herbivore microbiome with high plant biomass-degrading capacity. PLoS Genet 6.

2.	Callahan BJ, McMurdie PJ, Rosen MJ, Han AW, Johnson AJA, Holmes SP. 2016. DADA2: High-resolution sample inference from Illumina amplicon data. Nat Methods 13:581–583.

3.	Kohl KD, Dearing MD. 2012. Experience matters: Prior exposure to plant toxins enhances diversity of gut microbes in herbivores. Ecol Lett 15:1008–1015.

4.	Aylward FO, Suen G, Biedermann PHW, Adams AS, Scott JJ, Malfatti SA, Del Rio TG, Tringe SG, Poulsen M, Raffa KF, Klepzig KD, Currie CR. 2014. Convergent bacterial microbiotas in the fungal agricultural systems of insects. MBio 5:1–10.

5.	Aylward FO, Burnum KE, Scott JJ, Suen G, Tringe SG, Adams SM, Barry KW, Nicora CD, Piehowski PD, Purvine SO, Starrett GJ, Goodwin LA, Smith RD, Lipton MS, Currie CR. 2012. Metagenomic and metaproteomic insights into bacterial communities in leaf-cutter ant fungus gardens. ISME J 6:1688–1701.

6.	Khadempour L, Fan H, Keefover-Ring K, Carlos C, Nagamoto NS, Dam MA, Pupo MT, Currie CR. 2019. Metagenomics reveals diet-specific specialization of bacterial communities in fungus gardens of grass- and dicot-cutter ants. bioRxiv 250993.
