## Supplemental Table 1 for "Bacteria contribute to plant secondary compound degradation in a generalist herbivore system"

Table S1. Collection information for fungal isolates used in tolerance assay and bacterial isolates chosen for whole-genome sequencing.

| Isolate ID | Genus | Host colony | Host-ID | GPS Coordinates | Location | location notes | Date collected (m/d/y) | colony code |
|---|---|---|---|---|---|---|---|---|
| AS1 | Leucoagaricus | Atta sexdens | | -21.16337, -47.8615 | Sao Paulo, Brazil | | 11/27/14 | LK141127-01 |
| AB1 | Leucoagaricus | Atta bisphaerica | | -22.84652, -48.43411 | Sao Paulo, Brazil | | 11/21/15 | LK151121-01 |
| AC1 | Leucoagaricus | Atta capiguara | | -22.84642, -48.43427 | Sao Paulo, Brazil | | 11/21/15 | LK151121-05 |
| AL1 | Leucoagaricus | Atta laevigata | | -21.1675, -47.84638 | Sao Paulo, Brazil | | 12/5/15 | LK151205-01 |
| WM170124-07 | Leucoagaricus | Trachymyrmex diversus | 2255 | S2° 16' 13.7 W61° 01' 05.3 | Anavilhanas, AM | in fallen log | 1/24/17 | WM170124-07 |
| ICBG809 | Acinetobacter | Trachymyrmex | 2318 | S2° 55.825 W59° 58.529 | Ducke Reserve, AM | on coconut | 1/28/17 | WM170128-03 |
| ICBG1742 | Asaia | Atta sexdens | 2338 | -22.84652, -48.43411 | Sao Paulo State | Botucatu | 1/6/17 | LK170106-08 |
| ICBG1751 | Bacillus | Atta bisphaerica | 2342 | -22.84652, -48.43413 | Sao Paulo State | Botucatu | 1/6/17 | LK170106-01 |
| ICBG637 | Burkholderia | Cyphomyrmex | 2324 | S2° 55.808 W59° 58.479 | Ducke Reserve, AM | on log | 1/28/17 | TF170128-03 |
| ICBG647 | Burkholderia | Cyphomyrmex | 2320 | S2° 55.808 W59° 58.479 | Ducke Reserve, AM | on log | 1/28/17 | WM170128-05 |
| ICBG1735 | Burkholderia | Atta sp. | 2341 | -22.84652, -48.43412 | Sao Paulo State | Botucatu | 1/6/17 | LK170106-06 |
| ICBG862 | Burkholderia | Trachymyrmex | 2256 | S2° 16' 13.7 W61° 01' 05.3 | Anavilhanas, AM | in fallen log | 1/24/17 | WM170124-08 |
| ICBG955 | Burkholderia | Apterostigma | 2086 | S2° 32' 1.9 W60° 50' 6.1 | Anavilhanas, AM | under log | 1/19/17 | BP170119-02 |
| ICBG962 | Burkholderia | Apterostigma | 2149 | S2° 31' 23,4 W60° 49' 31,9 | Anavilhanas, AM | in fallen log | 1/20/17 | WM170120-10 |
| ICBG1719 | Burkholderia | Atta sp. | 2341 | -22.84652, -48.43412 | Sao Paulo State | Botucatu | 1/6/17 | LK170106-06 |
| ICBG1720 | Burkholderia | Atta sp. | 2341 | -22.84652, -48.43412 | Sao Paulo State | Botucatu | 1/6/17 | LK170106-06 |
| ICBG1724 | Burkholderia | Atta sp. | 2340 | -22.84652, -48.43411 | Sao Paulo State | Botucatu | 1/6/17 | LK170106-05 |
| ICBG849 | Burkholderia | Trachymyrmex | 2185 | S2° 34.818 W61° 02.045 | Anavilhanas, AM | | 1/23/17 | AR170123-03 |
| ICBG860 | Burkholderia | Trachymyrmex | 2256 | S2° 16' 13.7 W61° 01' 05.3 | Anavilhanas, AM | in fallen log | 1/24/17 | WM170124-08 |
| ICBG641 | Paraburkholderia | Cyphomyrmex | 2324 | S2° 55.808 W59° 58.479 | Ducke Reserve, AM | on log | 1/28/17 | TF170128-03 |
| ICBG1792 | Chitinophaga | Cyphomyrmex | 2186 | S2° 35.9 W61° 1.50 | Anavilhanas, AM | deep in large stum | 1/23/17 | LK170123-01 |
| ICBG659 | Chryseobacterium | Trachymyrmex | 2270 | n/a | Anavilhanas, AM | | 1/24/17 | CC170124-12 |
| ICBG654 | Comamonas | Trachymyrmex | 2270 | n/a | Anavilhanas, AM | | 1/24/17 | CC170124-12 |
| ICBG1797 | Enterobacter | Cyphomyrmex | 2186 | S2° 35.9 W61° 1.50 | Anavilhanas, AM | deep in large stum | 1/23/17 | LK170123-01 |
| ICBG867 | Enterobacter | Apterostigma | 2253 | S2° 16' 13.7 W61° 01' 05.3 | Anavilhanas, AM | beside tree | 1/24/17 | WM170124-05 |
| ICBG916 | Enterobacter | Apterostigma | 2311 | n/a | Ducke Reserve, AM | | 1/28/17 | CC170128-12 |
| ICBG1006 | Enterobacter | Apterostigma | 2253 | S2° 16' 13.7 W61° 01' 05.3 | Anavilhanas, AM | beside tree | 1/24/17 | WM170124-05 |
| ICBG643 | Enterobacter | Cyphomyrmex | 2324 | S2° 55.808 W59° 58.479 | Ducke Reserve, AM | on log | 1/28/17 | TF170128-03 |
| ICBG810 | Enterobacter | Trachymyrmex | 2318 | S2° 55.825 W59° 58.529 | Ducke Reserve, AM | on coconut | 1/28/17 | WM170128-03 |
| ICBG832 | Enterobacter | Acromyrmex | 2226 | S2° 16.262 W61° 01.141 | Anavilhanas, AM | | 1/24/17 | AR170124-01 |
| ICBG833 | Klebsiella | Acromyrmex | 2226 | S2° 16.262 W61° 01.141 | Anavilhanas, AM | | 1/24/17 | AR170124-01 |
| ICBG834 | Klebsiella | Acromyrmex | 2226 | S2° 16.262 W61° 01.141 | Anavilhanas, AM | | 1/24/17 | AR170124-01 |
| ICBG873 | Klebsiella | Apterostigma | 2253 | S2° 16' 13.7 W61° 01' 05.3 | Anavilhanas, AM | beside tree | 1/24/17 | WM170124-05 |
| ICBG874 | Klebsiella | Apterostigma | 2253 | S2° 16' 13.7 W61° 01' 05.3 | Anavilhanas, AM | beside tree | 1/24/17 | WM170124-05 |
| ICBG640 | Pantoea | Cyphomyrmex | 2324 | S2° 55.808 W59° 58.479 | Ducke Reserve, AM | on log | 1/28/17 | TF170128-03 |
| ICBG1710 | Pantoea | Atta laevigata | 2335 | 21°10'3"S 47°50'47"W | Sao Paulo State | Riberao Preto | 1/8/17 | LK170108-02 |
| ICBG1758 | Pantoea | Acromyrmex | 2334 | n/a | Sao Paulo State | Riberao Preto | 1/10/17 | CAR170110-02 |
| ICBG805 | Pantoea | Acromyrmex | 2160 | n/a | Anavilhanas, AM | | 1/21/17 | CAR170121-01 |
| ICBG807 | Pantoea | Acromyrmex | 2160 | n/a | Anavilhanas, AM | | 1/21/17 | CAR170121-01 |
| ICBG828 | Pantoea | Trachymyrmex | 2318 | S2° 55.825 W59° 58.529 | Ducke Reserve, AM | on coconut | 1/28/17 | WM170128-03 |
| ICBG835 | Pantoea | Acromyrmex | 2226 | S2° 16.262 W61° 01.141 | Anavilhanas, AM | | 1/24/17 | AR170124-01 |
| ICBG870 | Pantoea | Apterostigma | 2253 | S2° 16' 13.7 W61° 01' 05.3 | Anavilhanas, AM | beside tree | 1/24/17 | WM170124-05 |
| ICBG959 | Pantoea | Apterostigma | 2149 | S2° 31' 23,4 W60° 49' 31,9 | Anavilhanas, AM | in fallen log | 1/20/17 | WM170120-10 |
| ICBG985 | Pantoea | Apterostigma | 2311 | n/a | Ducke Reserve, AM | | 1/28/17 | CC170128-12 |
| ICBG639 | Pseudomonas | Cyphomyrmex | 2324 | S2° 55.808 W59° 58.479 | Ducke Reserve, AM | on log | 1/28/17 | TF170128-03 |
| ICBG811 | Pseudomonas | Trachymyrmex | 2318 | S2° 55.825 W59° 58.529 | Ducke Reserve, AM | on coconut | 1/28/17 | WM170128-03 |
| ICBG967 | Pseudomonas | Apterostigma | 2086 | S2° 32' 1.9 W60° 50' 6.1 | Anavilhanas, AM | under log | 1/19/17 | BP170119-02 |
