## Supplemental Table 2 for "Bacteria contribute to plant secondary compound degradation in a generalist herbivore system"

Table S2. Genome statistics and accession numbers for 42 bacterial isolates.

| Organism (16S identification) | ID number | Ant Host | Est. size (bp) | %GC | CDS | contigs | N50 | average coverage | Species (ANIm percent identity) | JspeciesWS tetra correlation search top hit | BioProject | Whole Genome Accession | SRA Accession |
|---|---|---|---|---|---|---|---|---|---|---|---|---|---|
| Asaia | ICBG1742 | Atta sexdens | 3744109 | 59.55 | 3369 | 70 | 455179 | 153 | A. bogorensis NBRC 16594 (.95) | A. sp. SF2.1 (.9972) | PRJNA564151 | VWWA00000000 | SRR10136620 |
| Burkholderia | ICBG862 | Trachymyrmex | 7986442 | 67.96 | 7028 | 958 | 16317 | 30 | B. gladioli ATCC 10248 (.97) | B. gladioli (GCA_001527485) MSMB1756 (.9996) | PRJNA564151 | VWWG00000000 | SRR10136585 |
| Burkholderia | ICBG637 | Cyphomyrmex | 7912056 | 68.26 | 6903 | 848 | 17648 | 27 | B. gladioli ATCC 10248 (.97) | B. gladioli (GCA_001527485) MSMB1756 (.9997) | PRJNA564151 | VWWC00000000 | SRR10136598 |
| Burkholderia | ICBG647 | Cyphomyrmex | 8207449 | 68.13 | 7053 | 482 | 34978 | 51 | B. gladioli ATCC 10248 (.97) | B. gladioli (GCA_001527485) MSMB1756 (.9997) | PRJNA564151 | VWWD00000000 | SRR10136588 |
| Burkholderia | ICBG955 | Apterostigma | 7982062 | 68.29 | 6819 | 422 | 36080 | 38 | B. gladioli ATCC 10248 (.97) | B. gladioli (GCA_001527485) MSMB1756 (.9995) | PRJNA564151 | VWWH00000000 | SRR10136584 |
| Burkholderia | ICBG962 | Apterostigma | 7913770 | 67.74 | 7111 | 1225 | 12004 | 28 | B. gladioli ATCC 10248 (.97) | B. gladioli (GCA_001527485) MSMB1756 (.9991) | PRJNA564151 | VWWI00000000 | SRR10136583 |
| Burkholderia | ICBG1735 | Atta sp. | 8136338 | 66.59 | 7226 | 240 | 67746 | 39 | B. lata FL-7-5-30-S1-D0 (.98) | B. lata (GCA_001718575) FL-7-5-30-S1-D0 (.9997) | PRJNA564151 | VWWE00000000 | SRR10136587 |
| Burkholderia | ICBG1719 | Atta sp. | 8174181 | 66.41 | 7336 | 472 | 44435 | 49 | B. lata FL-7-5-30-S1-D0 (.98) | B. lata (GCA_001718575) FL-7-5-30-S1-D0 (.9979) | PRJNA564151 | VWWJ00000000 | SRR10136619 |
| Burkholderia | ICBG1720 | Atta sp. | 8137953 | 66.47 | 7287 | 347 | 52454 | 42 | B. lata FL-7-5-30-S1-D0 (.98) | B. lata (GCA_001718575) FL-7-5-30-S1-D0 (.9991) | PRJNA564151 | VWWK00000000 | SRR10136618 |
| Burkholderia | ICBG1724 | Atta sp. | 7497824 | 66.3 | 6781 | 423 | 40856 | 37 | B. ambifaria AMMD (.96) | B. ambifaria MEX-5 (.9977) | PRJNA564151 | VWWL00000000 | SRR10136617 |
| Burkholderia | ICBG849 | Trachymyrmex | 7232450 | 63.74 | 6330 | 75 | 212914 | 52 | B. sp. H160 (.89) | B. sp. H160 (.987) | PRJNA564151 | VWWF00000000 | SRR10136586 |
| Burkholderia | ICBG860 | Trachymyrmex | 7049452 | 63.7 | 6208 | 140 | 109899 | 41 | B. sp. H160 (.89) | B. sp. H160 (.9871) | PRJNA564151 | VWWN00000000 | SRR10136615 |
| Burkholderia | ICBG641 | Cyphomyrmex | 8127228 | 62.99 | 7066 | 78 | 264938 | 26 | Paraburkholderia eburnea (.89) | Paraburkholderia eburnea JCM 18070 (.9799) | PRJNA564151 | VWWM00000000 | SRR10136616 |
| Comamonas | ICBG654 | Trachymyrmex | 5460670 | 61.81 | 4815 | 29 | 812373 | 74 | C. testosteroni TK102 (.96) | C. testosteroni NBRC 14951 (.9986) | PRJNA564151 | VWWQ00000000 | SRR10136612 |
| Acinetobacter | ICBG809 | Trachymyrmex | 3864618 | 40.57 | 3608 | 50 | 512398 | 81 | A. gyllenbergii NIPH230 (.88) | A. sp. NIPH 1867 (.9965) | PRJNA564151 | VWVZ00000000 | SRR10136621 |
| Cedecea | ICBG1797 | Cyphomyrmex | 4783738 | 54.84 | 4425 | 26 | 2804548 | 70 | E. sp. Ag1 (.948) | E. sp. Ag1 (.9991) | PRJNA564151 | VWWR00000000 | SRR10136611 |
| Cedecea | ICBG916 | Apterostigma | 4734509 | 54.95 | 4345 | 36 | 1179865 | 64 | E.sp. Ag1 (.979) | E. sp. Ag1 (.9996) | PRJNA564151 | VWWT00000000 | SRR10136608 |
| Cedecea | ICBG1006 | Apterostigma | 4763828 | 54.87 | 4400 | 36 | 497031 | 106 | E. sp. Ag1 (.989) | E. sp. Ag1 (.9998) | PRJNA564151 | VWWU00000000 | SRR10136607 |
| Cedecea | ICBG867 | Apterostigma | 4782879 | 54.87 | 4424 | 52 | 781065 | 72 | E. sp. Ag1 (.989) | E. sp. Ag1 (.9998) | PRJNA564151 | VWWS00000000 | SRR10136610 |
| Enterobacter | ICBG832 | Acromyrmex | 4490220 | 55.66 | 4134 | 46 | 2412389 | 92 | E. sp. CRENT-193 (.99) | E. cloacae (GCA_001375695) Marseille (.9998) | PRJNA564151 | VWWX00000000 | SRR10136604 |
| Enterobacter | ICBG643 | Cyphomyrmex | 4756147 | 53.86 | 4380 | 32 | 562458 | 57 | E. asburiae LF7a (.98) | E. soli ATCC BAA-2102 (.9996) | PRJNA564151 | VWWV00000000 | SRR10136606 |
| Enterobacter | ICBG810 | Trachymyrmex | 4858342 | 55.87 | 4498 | 35 | 712084 | 81 | E. cloacae DSM16690 (.99) | E. sp. ku-bf2 (.9996) | PRJNA564151 | VWWW00000000 | SRR10136605 |
| Pantoea | ICBG807 | Acromyrmex | 5859404 | 53.67 | 5555 | 98 | 233234 | 47 | P. sp. GL120224-02 (.97) | P. sp. BL1 (.999) | PRJNA564151 | VWXE00000000 | SRR10136595 |
| Pantoea | ICBG805 | Acromyrmex | 5056091 | 57.32 | 4688 | 39 | 478141 | 113 | P. sp. VS1 (.98) | P. dispersa SA5 (.9995) | PRJNA564151 | VWXD00000000 | SRR10136596 |
| Pantoea | ICBG1758 | Acromrymex | 4177385 | 56.17 | 3855 | 47 | 479536 | 121 | P. sp. Ae16 (.99) | P. eucrina LMG 5346 (.9995) | PRJNA429666 | POWL00000000 | SRR10145087 |
| Pantoea | ICBG835 | Acromyrmex | 5903156 | 52.91 | 5530 | 84 | 695466 | 75 | P. rwandensis LMG 26275 (.92) | P. rwandensis LMG 26275 (.9979) | PRJNA564151 | VWXF00000000 | SRR10136594 |
| Pantoea | ICBG870 | Apterostigma | 5421039 | 52.83 | 4857 | 56 | 418101 | 95 | P. sp A4 (.96) | P. rodasii ND03 (.9681) | PRJNA564151 | VWXG00000000 | SRR10136593 |
| Pantoea | ICBG959 | Apterostigma | 4990646 | 57.41 | 4519 | 68 | 460846 | 96 | P. dispersa EGD AAK13 (.98) | P. dispersa SA5 (.9996) | PRJNA564151 | VWXH00000000 | SRR10136592 |
| Pantoea | ICBG985 | Apterostigma | 4932224 | 57.36 | 4509 | 37 | 703296 | 66 | P. dispersa SA5 (.98) | P. dispersa SA5 (.9996) | PRJNA429667 | POWM00000000 | SRR10145031 |
| Pantoea | ICBG1710 | Atta laevigata | 3983324 | 56.54 | 3635 | 54 | 459904 | 108 | P. sp. Ae16 (.99) | P. eucrina LMG 5346 (.9996) | PRJNA564151 | VWXC00000000 | SRR10136597 |
| Pantoea | ICBG640 | Cyphomyrmex | 5027706 | 57.52 | 4568 | 54 | 613347 | 53 | P. dispersa EGD AAK13 (.98) | P. dispersa SA5 (.9997) | PRJNA564151 | VWXB00000000 | SRR10136599 |
| Pantoea | ICBG828 | Trachymyrmex | 5026145 | 57.52 | 4565 | 51 | 551420 | 73 | P. dispersa EGD AAK13 (.98) | P. dispersa SA5 (.9997) | PRJNA429668 | POWN00000000 | SRR10145062 |
| Klebsiella | ICBG833 | Acromyrmex | 5718030 | 57.24 | 5307 | 50 | 772383 | 64 | K. variicola GJ1 (.99) | K. pneumoniae MGH 80 (.9999) | PRJNA564151 | VWWY00000000 | SRR10136603 |
| Klebsiella | ICBG834 | Acromyrmex | 5719244 | 57.24 | 5305 | 54 | 772383 | 59 | K. variicola GJ1 (.99) | K. pneumoniae MGH 80 (.9999) | PRJNA564151 | VWWZ00000000 | SRR10136602 |
| Klebsiella | ICBG873 | Apterostigma | 10438772 | 55.95 | 9773 | 91 | 617715 | 37 | K. variicola At-22 (.98) | K. pneumoniae IS22 (.9886) | PRJNA564151 | JAADCL000000000 | SRR10136601 |
| Klebsiella | ICBG874 | Apterostigma | 5395446 | 57.53 | 4987 | 45 | 954810 | 82 | K. variicola At-22 (.99) | K. variicola (GCA_001261875) (.9999) | PRJNA564151 | VWXA00000000 | SRR10136600 |
| Pseudomonas | ICBG967 | Apterostigma | 5262567 | 64.51 | 4597 | 78 | 233438 | 50 | P. mosselii BS011 (.89) | P. soli LMG 27941 (.9949) | PRJNA564151 | VWXK00000000 | SRR10136589 |
| Pseudomonas | ICBG639 | Cyphomyrmex | 6659621 | 63.5 | 5755 | 109 | 297835 | 49 | P. putida AA7 (.89) | P. sp. GM84 (.9862) | PRJNA564151 | VWXI00000000 | SRR10136591 |
| Pseudomonas | ICBG811 | Trachymyrmex | 6672247 | 63.5 | 5806 | 142 | 192183 | 49 | P. putida AA7 (.89) | P. sp. GM84 (.9865) | PRJNA564151 | VWXJ00000000 | SRR10136590 |
| Chitinophaga | ICBG1792 | Cyphomyrmex | 7569413 | 45.59 | 6072 | 51 | 2808195 | 65 | C. bacterium IBVUCB2 (.87) | C. jiangningensis DSM 27406 (.9755) | PRJNA564151 | VWWO00000000 | SRR10136614 |
| Chryseobacterium | ICBG659 | Trachymyrmex | 4879137 | 36.18 | 4264 | 39 | 354653 | 119 | C. sp. CF365 (.86) | C. sp. GSE06 (.9861) | PRJNA564151 | VWWP00000000 | SRR10136613 |
| Bacillus | ICBG1751 | Atta bisphaerica | 5884491 | 34.77 | 5795 | 67 | 459078 | 67 | B. thuringiensis YBT-1518 (.96) | B. thuringiensis (GCA_001757745) GOE4 (.9997) | PRJNA564151 | VWWB00000000 | SRR10136609 |
