## Supplemental Table 3 for "Bacteria contribute to plant secondary compound degradation in a generalist herbivore system"

Table S3. Laboratory colony collection information used for headspace sampling and 16S rRNA amplicon experiments.

| Colony Name | Colony Code | Ant species | GPS | Location | Country |
|---|---|---|---|---|---|
| Dora | RM120223-02 | Atta cephalotes | n/a | La Selva Biological Station | Costa Rica |
| Regina | CR14 | Atta cephalotes | n/a | Finca la Anita | Costa Rica |
| Leslie | CF180406-01 | Atta cephalotes | N10.42764, W84.00176 | La Selva Biological Station | Costa Rica |
| Joan | CF180405-02 | Atta cephalotes | N10.42338, W84.00136 | La Selva Biological Station | Costa Rica |
| Mona-Lisa | HH180403-03 | Atta cephalotes | N10.43136, W84.00557 | La Selva Biological Station | Costa Rica |
