## Supplemental Table 4 for "Bacteria contribute to plant secondary compound degradation in a generalist herbivore system"

Table S4. Fungus garden collection information for metatranscriptomic sequencing.

| ID | Collection Date | Uploaded to MG-RAST | Ant species | Location | Country | Mean Sequence Length (post-QC) | BioProject | SRA Accession |
|---|---|---|---|---|---|---|---|---|
| FG1 | June 1 2014 | 2/26/15 | Atta cephalotes | La Selva Biological Station | Costa Rica | 148 ± 24 bp | PRJNA565936 | SRR10132760 |
| FG2 | June 1 2014 | 2/26/15 | Atta cephalotes | La Selva Biological Station | Costa Rica | 143 ± 22 bp | PRJNA565936 | SRR10132759 |
| FG3 | Feb 25 2014 | 2/26/15 | Atta colombica | Golfito | Costa Rica | 145 ± 23 bp | PRJNA565936 | SRR10132758 |
